# Hydrophobic tuning with non-canonical amino acids in a copper metalloenzyme

**DOI:** 10.1101/2025.02.09.636911

**Authors:** Sandro Fischer, Anton Natter Perdiguero, Kelvin Lau, Alexandria Deliz Liang

## Abstract

Hydrophobicity controls many aspects of protein and enzyme function. Although hydrophobic tuning can be achieved to a limited extent with canonical amino acids, the incorporation of non-canonical amino acids (ncAAs) further extends this ability to enable new and improved functionality. Herein, we engineer an aminoacyl-tRNA synthetase/tRNA pair for the site-specific genetic encoding of a set of bulky, hydrophobic amino acids, namely cyclopentylalanine, cyclohexylalanine, and cycloheptylalanine. With the resulting orthogonal translations systems, we demonstrate the utility of ncAA-based hydrophobic tuning to engineer a bacterial laccase, which is both a classical metalloenzyme and a high-value catalyst for industrial processes. The resulting mutations conveyed significant improvements in catalytic activity, particularly *k*_cat_ and total turnover number. The redox potential and structure-function relationships were examined to elucidate the source of this improved functionality. We envision that these tools for hydrophobic tuning will be highly valuable for general enzyme engineering and also in other fields, including peptide chemistry.

## Introduction

Hydrophobic interactions are crucial for stabilizing protein cores and driving folding processes. Within enzymes, hydrophobic environments also play important roles in promoting catalysis, tuning cofactor binding, substrate binding, and reactivity^1,2^. In the laboratory, modifications of such environments by programmed mutagenesis have demonstrated both chemically intuitive and surprising effects in proteins and enzymes. For metalloenzymes in particular, hydrophobic environments can affect a plethora of properties, including the redox potential of redox-active metal ions^3^, the pK_a_ of metal-coordinated water molecules^4^, and even metal specificity^5,6^. Both nature and traditional laboratory mutagenesis use predominantly 20 canonical amino acids to optimize or study these effects. However, using only canonical amino acids limits the possibilities for hydrophobic tuning. In contrast, incorporation of non-canonical amino acids (ncAAs) allows for broadened chemical variability, enabling detailed enzymatic studies^7–10^, novel reactions^11–14^, and even improved function^15–18^. A few studies have demonstrated improved enzyme and protein function using hydrophobic tuning with proteins generated through native chemical ligation^3,19^, in vitro protein production with pre-aminoacylated tRNAs^20^, and global ncAA replacement of methionine with norleucine or homopropargylglycine^21–23^. However, these techniques are limited compared to the genetic incorporation of ncAAs in a site-specific manner. Such limitations include low protein yield, limited applicability to diverse protein scaffolds, and a lack of targeted mutagenesis, preventing (semi-)rational mutagenesis.

Despite the value of site-specific, genetic ncAA incorporation, this strategy remains underexplored for hydrophobic tuning, largely owing to the limited available tools. Herein, we address this challenge by developing a platform for the robust genetic incorporation of three bulky, hydrophobic ncAAs: L-3-cyclopentylalanine (**C5a**), L-3-cyclohexylalanine (**C6a**), and L-3-cycloheptylalanine (**C7a**). These novel tools were then applied to fine-tune the hydrophobic environment of a high-value copper enzyme—a laccase (EC 1.10.3.2) with a prototypical monocopper center.

Laccases oxidize diverse substrates concomitantly reducing oxygen to water^24,25^, making them “green” catalysts for degradation of pollutants, biomass valorization^26,27^, product bleaching^28^, synthetic chemistry^29,30^, bioremediation^31^, food treatment^28^, and textile processing^32^. Laccases are categorized as low- or high-redox potential enzymes according to the redox potential of their T1Cu center (E°’_T1Cu_), which ranges between 360 and 785 mV vs the normal hydrogen electrode (NHE)^33^. High-redox potential laccases—typically found in fungi—are potent catalysts, but display reduced stability, require complex glycosylation patterns for function, often require long expression times with low yields, and are not suitable for production in *E. coli*^34–36^. In contrast, low-redox laccases—typically found in bacteria—can be rapidly produced with high yield and are active under harsh conditions^36^. Unfortunately, low-redox laccases are less applicable in many domains because of their diminished activity^34^, which is often linked to their low redox potential. Several attempts have been made to increase the redox potential of bacterial laccases by engineering the primary coordination sphere of the T1Cu site to resemble the T1Cu site of high-redox potential laccases^37–43^. These classical mutagenesis methods have been shown to increase the redox potential for some enzymes, but significantly diminish catalytic activity in all cases^39,41,44–48^. These studies illustrate a major challenge in metal-enzyme engineering: the concomitant optimization of electronic properties, substrate binding, protein stability, and catalytic efficiency.

We posited that that hydrophobic tuning of the bacterial laccase from *Streptomyces coelicolor* (*Sc*SLAC) could increase the redox potential, improve catalysis, and overcome prior limitations encountered when engineering the primary coordination sphere of the T1Cu environment. We demonstrate that site-selective incorporation of either **C5a** or **C6a** at a key position in the active site increases the redox potential and improves catalytic performance, particularly with phenolic small molecules that are often used as lignin surrogates. These results demonstrate that site-specific incorporation of ncAAs can be used to improve catalysis, tune a hydrophobic environment, and directly promote the rate-limiting step(s) of an enzyme.

## Results

### An engineered aaRS/tRNA pair for incorporation of hydrophobic ncAAs

We first identified three ncAAs that are hydrophobic, space-filling, and vary in size (Figure 1a): L-3-cyclopentylalanine (**C5a**), L-3-cyclohexylalanine (**C6a**), and L-3-cycloheptylalanine (**C7a**). Site-specific incorporation of ncAAs in cells requires the development of an aminoacyl-tRNA synthetase (aaRS)/tRNA pair that specifically recognizes an ncAA and enables recoding of a target codon via the ribosome. Typically, the codon of choice is the amber stop codon (TAG). Due to the nature of aaRS/tRNA engineering, the evolved aaRS/tRNA pair can often selectively incorporate the desired ncAA over canonical amino acids but retains the ability to incorporate ncAAs of similar structure. This feature, termed polyspecificity, is a powerful way to incorporate various ncAAs using a single aaRS/tRNA, enabling facile ncAA mutagenesis. To leverage polyspecificity, we engineered an aaRS/tRNA pair for incorporation of **C6a**, which is intermediate in size between the three selected ncAAs, and thus would give us the best chance of obtaining a polyspecific aaRS/tRNA pair. The pyrrolysyl-tRNA synthetase (PylRS)/tRNA pair was selected as the parent system for further engineering based on its evolvability and its orthogonality in several common laboratory hosts^49^. A set of libraries of PylRS from *Methanosarcina mazei* (*Mm*PylRS) was prepared by randomizing between three and five residues of the substrate binding site using site-saturation mutagenesis (Figure 1b). Additionally, a library of *Mm*PylRS containing the fixed I405R mutation was prepared^50^. The libraries were pooled and subjected to a round of positive selection in the presence of 6 mM **C6a** using an sfGFP reporter containing an amber stop codon at position N150 (sfGFP150_TAG_). Individual clones were screened for ncAA selectivity by evaluating amber suppression in the presence and absence of 6 mM **C6a**. An *Mm*PylRS variant (PylRS^C6av0^) was identified that exhibited an sfGFP-derived fluorescence increase of approximately 2.6-fold upon addition of 6 mM **C6a** (Supplementary Figure 1a). To improve the orthogonality of PylRS^C6av0^, additional sites were selected for a second round of site-saturation mutagenesis, and the libraries were subjected to rounds of alternating positive and negative selection. The best variant, termed PylRS^C6a^, displayed reduced suppression in the absence of **C6a** and an approximately four-fold fluorescence increase in suppression upon addition of 6 mM **C6a** (Supplementary Figure 1b).

**Figure 1.**
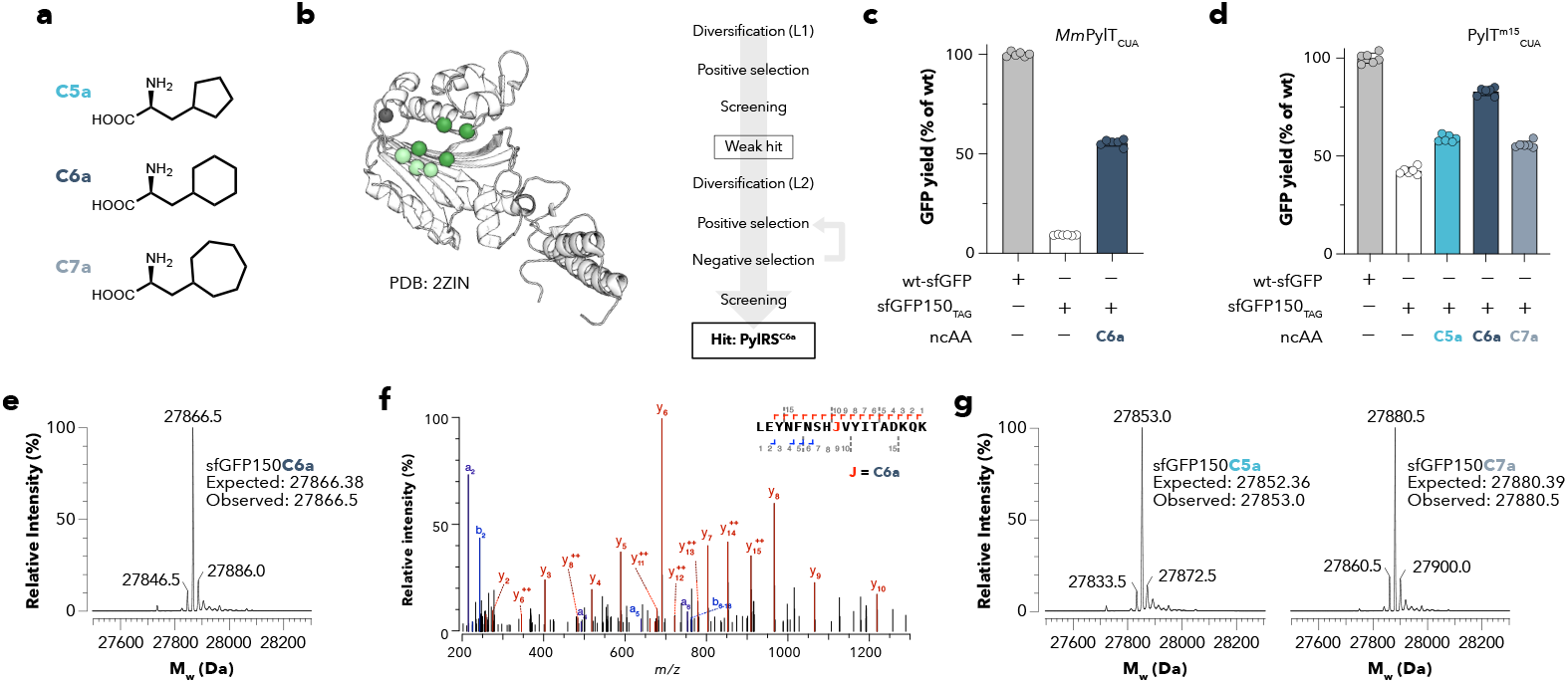
Engineering an aaRS/tRNA pair for bulky hydrophobic ncAAs. a) Structures of ncAAs incorporated in this study: L-3-cyclopentylalanine (**C5a**), L-3-cyclohexylalanine (**C6a**), and L-3-cycloheptylalanine (**C7a**). b) Mutation positions within libraries used for engineering PylRS^C6a^ from *Mm*PylRS, PDB 2ZIN.^51^ c) Amber suppression benchmarking of pGS1T-PylRS^C6a^-*Mm*PylT_CUA_ pair in sfGFP150_TAG_ with and without 6 mM **C6a**. d) amber suppression benchmarking of pGS1T-PylRS^C6a^-PylT^m15^_CUA_ pair in sfGFP150_TAG_ with and without 12 mM DL-**C5a**, 6 mM **C6a**, or 12 mM DL-**C7a**. Data are normalized to an otherwise wild-type protein (wt sfGFP) and represent the mean and standard deviation of six biological replicates. e) Intact LC-MS analysis (positive electrospray time of flight) of sfGFP150_TAG_ expression in the presence of 6 mM **C6a**. Additional peaks consistent with typically observed [H_2_O] elimination (−18 Da) and [Na]^+^ addition (+23 Da) are labeled. f) A representative LC-MS/MS spectrum from a tryptic digest of sfGFP150_TAG_ expressed with pGS1T-PylRSC6a-PylT^m15^_CUA_ in the presence of 6 mM **C6a**. Multiple peptides containing **C6a** were observed, but no peptides for canonical amino acid incorporation were observed. g) Intact LC-MS analysis (positive electrospray time of flight) of sfGFP150_TAG_ expression in the presence of 12 mM DL-**C5a** or 12 mM DL-**C7a**. Additional peaks consistent with typically observed [H_2_O] elimination (−18 Da) and [Na]^+^ addition (+23 Da) are labeled. The complete collected MS spectra for insets e and g are supplied in Supplementary Figure 2.

We designed an optimized plasmid system for the expression of engineered PylRS/tRNA pairs (pGS1T). The PylRS^C6a^/*Mm*PylT_CUA_ pair was cloned onto pGS1T where PylRS^C6a^ is under the control of a constitutive glnS promoter and *Mm*PylT_CUA_ under a proK promoter. In the optimized plasmid system, expression of sfGFP150_TAG_ from a pBAD vector in NEB10β showed an approximately six-fold increase upon addition of 6 mM **C6a**, yielding >50% of wild-type protein (wt sfGFP) production (Figure 1c). To further improve the suppression efficiency, *Mm*PylT_CUA_ was replaced with PylT^m15^_CUA_^52^. The resulting construct improved the incorporation efficiency of **C6a** in sfGFP to result in wt-like levels of sfGFP production in the presence of 6 mM **C6a** (Figure 1d). In the absence of **C6a**, incorporation of phenylalanine was observed. LC-MS analysis of intact sfGFP150_TAG_ obtained in the presence of **C6a** showed incorporation of only the desired species (Figure 1e). Importantly, no peaks corresponding to incorporation of any other canonical amino acid could be observed, indicating selectivity in the presence of **C6a**. To further rule out incorporation of canonical amino acids, we additionally confirmed the selective incorporation of **C6a** by trypsin digestion and LC-MS/MS (Figure 1f). Although multiple peptides containing **C6a** were observed, no peptides for canonical amino acids were observed. Lastly, we tested the polyspecificity of the evolved PylRS^C6a^ towards the two additional ncAAs **C5a** and **C7a**. We expressed sfGFP150_TAG_ in the presence and absence of the ncAAs and observed a slight increase in sfGFP production upon addition of 12 mM DL-ncAA for both ncAAs (Figure 1d). In light of the high phenylalanine incorporation in the absence of an ncAA, we performed LC-MS to determine the identity of the incorporated ncAA. LC-MS data indicated incorporation of the desired ncAAs without phenylalanine incorporation (Figure 1g). Thus, we conclude that in the presence of the desired ncAA, the engineered PylRS^C6a^/PylT^m15^_CUA_ is capable of selective incorporation of **C5a, C6a**, or **C7a** into proteins in *E. coli*. Alternatively, if selectivity in the absence of the ncAA is desired, the engineered PylRS^C6a^/*Mm*PylT_CUA_ pair is a highly suitable alternative.

### Replacement of axial methionine with C5a or C6a increases the redox potential

Laccases contain two distinct copper sites that are responsible for catalysis: a prototypical mononuclear T1Cu site and a trinuclear copper site, comprising a T2Cu and a T3Cu site (Figure 2a)^24^. The catalytic activity of laccases is dependent on the T1Cu site that oxidizes substrates. One of the notable differences between low- and high-redox potential laccases is the coordination at the T1Cu site. Low-redox potential laccases contain a CysHis_2_Met motif (Figure 2b). In contrast, high-redox potential laccases contain a CysHis_2_ motif, where the analogous methionine residue is replaced by a non-coordinating phenylalanine or leucine residue. In bacterial laccases, direct mutation of the methionine to phenylalanine or leucine has been shown to alter the redox potential^47,48^, but these fungal-like mutations also cause local structural perturbations and decrease in catalytic efficiency^39,41,44–46^.

**Figure 2.**
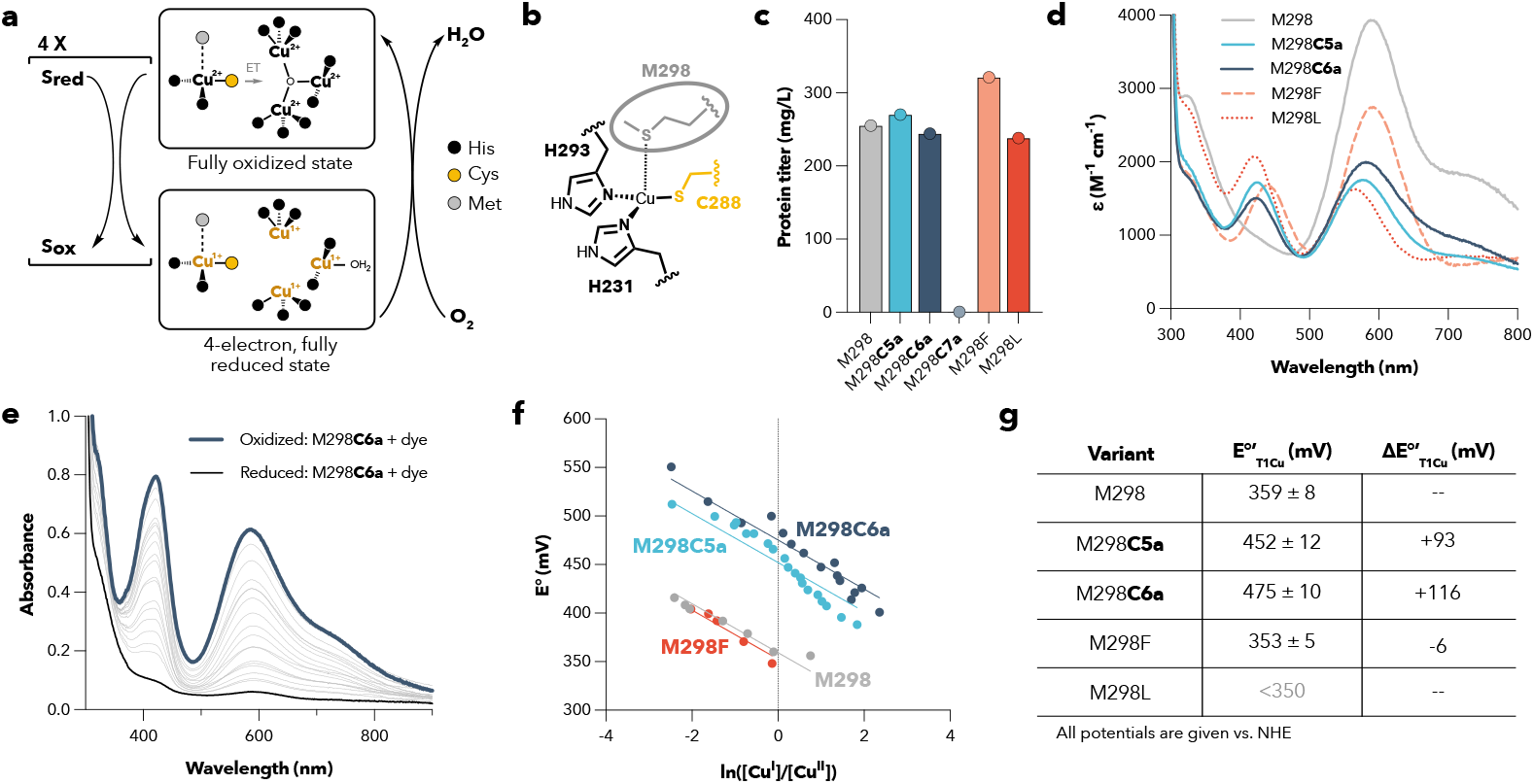
*Sc*SLAC: general reactivity, orientation of copper centers, and variant characterization: M298, M298C5a, M298C6a, M298C7a, M298F, and M298L. a) General substrate oxidation and oxidase activity of *Sc*SLAC. b) Coordination chemistry of the *Sc*SLAC T1Cu site. c) Expression titer of *Sc*SLAC variants in mg/L of culture from a single, representative, expression. d) Absorption spectra of *Sc*SLAC variants. e) Spectrophotometric redox titration of M298**C6a** with potassium ferricyanide as a redox indicator at pH 7.0. f) Analysis of the ratio of oxidized to reduced T1Cu species at varying potentials during the spectrophotometric titration. g) Estimated redox potentials for *Sc*SLAC variants based on the mean of the redox titration ± the standard deviation. The potential of M298L was too low to be determined by the spectrophotometric method.

We postulated that mutation of the axial methionine to the bulky ncAAs—**C5a, C6a**, or **C7a**— could lower the electron density at the T1Cu site, provide a more hydrophobic environment, and prevent water from accumulating in the T1Cu site; all of which could increase the redox potential and ideally improve catalysis. Thus, we designed experiments to incorporate the ncAAs **C5a, C6a**, and **C7a** into the axial position of the T1Cu site in *Sc*SLAC (M298, Figure 2b). We produced the desired *Sc*SLAC variants using the engineered PylRS^C6a^/PylT^m15^_CUA_ in the strain B-95ΔAΔfabR, which has been previously used for high-yielding, truncation-free genetic code expansion^53,54^. Gratifyingly, we could obtain high, wt-like titers for the two smaller ncAAs **C5a** and **C6a** (Figure 2c). However, no soluble protein was obtained for the larger **C7a** (Figure 2c). We postulate that the increased size of **C7a** precludes production of soluble *Sc*SLAC. Additionally, we isolated wt *Sc*SLAC and the high-redox potential mimics—mutants M298F and M298L—to compare the properties of these key mutations. For each protein variant, we conducted intact mass spectrometry (Supplementary Figure 3). All protein variants were found to have the expected mass.

In T1Cu-containing proteins, the ultraviolet-visible (UV-VIS) absorption spectra of the oxidized Cu^2+^ states are often used as descriptors for the copper center. In the visible region, two ligand-to-metal charge-transfer (LMCT) bands can occur, typically from a π-bonding interaction (∼600 nm) and a σ-bonding interaction (∼450 nm) between the sulfur p-orbitals of the cysteine to the 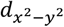 orbital of the copper ion^47,55^. The relative intensities of each of these bands (ε_σ/π_) have been connected to the extent of tetragonal distortion from a pseudo-tetrahedral His_2_CysMet T1Cu or exchange of the coordinating ligands^24^. Surprisingly, a continuum of relative intensities exists even for sites with the same coordination motifs. Such that both “blue” (low ε_σ/π_) and “green” (high ε_σ/π_) copper proteins can arise from the canonical His_2_CysMet T1Cu motif^24^. To compare the UV-VIS features of our *Sc*SLAC variants, we collected the UV-VIS absorption spectra of each oxidized variant. Compared to wt *Sc*SLAC, all mutants showed significant decrease in the intensity of the band around 600 nm and increase in intensity of the band around 450 nm (Figure 2d), which is reflected in their ε_σ/π_: wt *Sc*SLAC, 0.30; M298C5a, 0.98; M298C6a, 0.75; M298F, 0.61; and M298L, 1.28. These results are consistent with a modified coordination environment, but given the continuum of possible spectra for a given species, structural insight is required to understand how the coordination environment has changed (vide infra).

To further probe the electronic differences between each variant, we determined the redox potentials of the T1Cu site by redox titrations with potassium ferricyanide (K_3_[Fe(CN)_6_]) as a redox indicator at pH 7.0 (Figure 2e-g, Supplementary Figure 4). The redox potential of the wild-type enzyme was determined to be 359(7) mV vs NHE, which is in agreement with literature values between 360-430 mV^43,56–58^. *Sc*SLAC variants with ncAA mutations exhibited significantly increased redox potentials compared to wt *Sc*SLAC (Figure 2g): +93 mV for *Sc*SLAC-M298**C5a** and +116 mV for ScSLAC-M298**C6a**. Surprisingly, for the fungal-like *Sc*SLAC M298F and M298L mutants, the redox potential was reduced compared to the wt *S*cSLAC. The potential of M298L was so low that it could not be accurately determined with the potassium ferricyanide titration method. These results are consistent with a previous report of *Sc*SLAC^47^, but are different from that observed for CotA from *Bacillus subtilis*^45^ and the small laccase from *Streptomyces sviceus*^41^, which display increases in redox potential between 40-90 mV for their fungal-like mutants. Nonetheless, for all these previously reported, fungal-like variants, the *k*_cat_ of substrate oxidation was significantly diminished.

### Axial C5a and C6a mutations improve catalysis

Intrigued by the redox potential results, we characterized the catalytic activity of each variant (Figure 3). The common laccase substrate 2,2’-azino-bis(3-ethylbenzothiazoline-6-sulfonic acid) (ABTS, 680 mV vs NHE^59^) was used to benchmark the activity (Figure 3a), and Michaelis-Menten kinetics were determined at pH 4 (Figure 3a-c, Supplementary Figure 5, and Supplementary Table 1). The catalytic constants (*k*_*cat*_) of the two fungal-like *Sc*SLAC variants M298F and M298L were eight- and five-fold reduced from wt *Sc*SLAC, respectively. In contrast, the mutation of the axial ligand to M298**C5a** and M298**C6a** increased the *k*_*cat*_ by approximately two-fold for each variant. Additionally, the *k*_cat_/K_m_ also increased for M298**C5a** and M298**C6a** by 1.6 and 1.8-fold, respectively.

**Figure 3.**
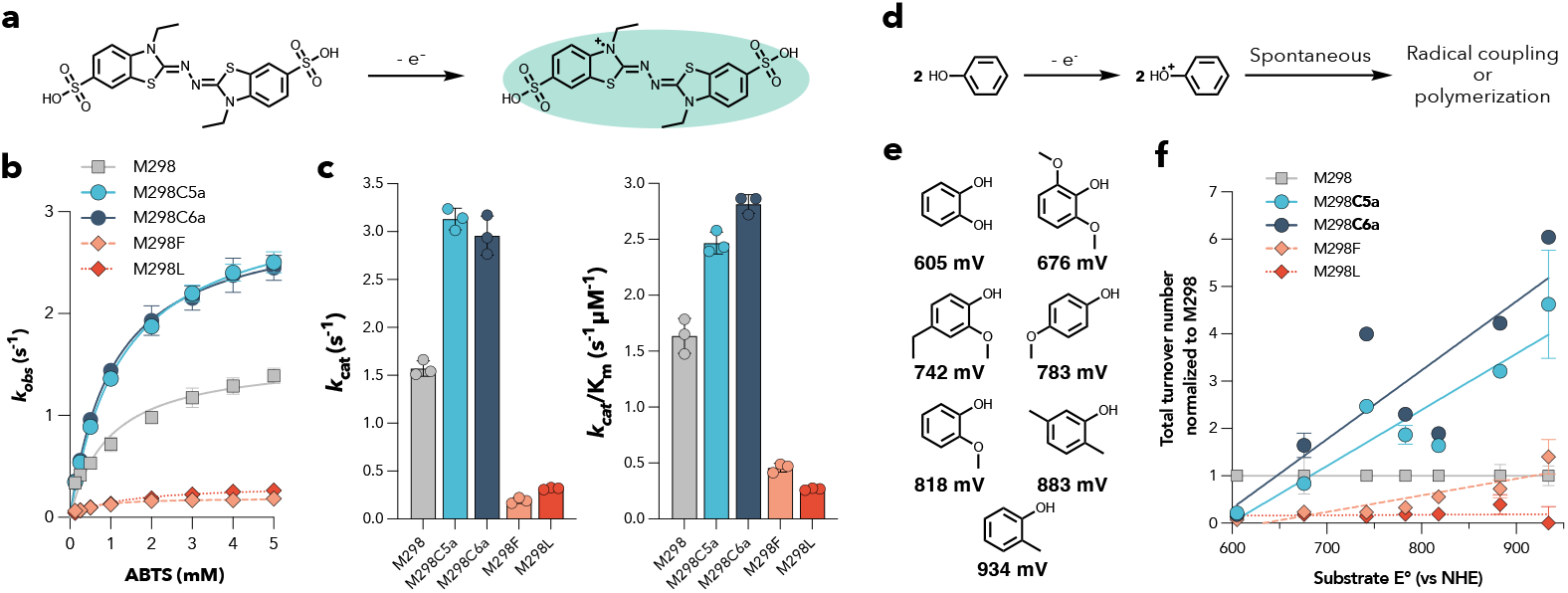
Characterization of steady-state activity and phenolic substrate profiling of *Sc*SLAC variants. a) Oxidation reaction of the classical laccase substrate ABTS. b) Michaelis-Menten kinetics of *Sc*SLAC variants with ABTS as a substrate at pH 4.0 at 25 °C. Data shown are biological triplicates. c) Comparison of kinetic parameters for the *Sc*SLAC variants. d) Phenolic oxidation reactions with laccases. e) Phenolic substrate panel and reported redox potentials vs. NHE. f) Total turnover numbers (TTN) of *Sc*SLAC variants with phenolic substrates at pH 7.0 at 25 °C. The TTNs are normalized to the TTN of *Sc*SLAC M298. Absolute values are shown in Supplementary Figure 6. Reactions were performed in biological triplicates. The error bars represent the standard deviation.

To assess the oxidation activity with higher redox potential substrates, the activity of *Sc*SLAC variants was assessed against a panel of phenolic compounds with varying redox potentials^60^ (Figure 3d-f, Supplementary Figures 6 and 7). For each variant, we determined the total turnover numbers (TTN) after 24 h. For M298**C5a** and M298**C6a**, a general trend was observed of increasing TTN relative to wt *Sc*SLAC. With the highest potential substrates, M298**C5a** and M298**C6a** showed an improvement of TTN compared to wt of up to five- and six-fold (Figure 3f). These results indicate that the ncAA-containing *Sc*SLAC variants can be used for oxidation of redox inert substrates and suggests the potential to oxidize molecules that are intractable with wt *Sc*SLAC. In contrast, the fungal-like variants M298F and M298L exhibit lower TTNs for almost all substrates compared to wt *Sc*SLAC.

Although a general trend is clear, we observed some outliers that are suggestive of some substrate-specific effects for M298**C5a** and M298**C6a**. These findings indicate that the activity cannot be explained by the T1Cu characteristics alone but is also dependent on substrate recognition. Notably, the activity is higher than expected with the substrate 4-ethyl-2-methoxyphenol (E° = 742 mV vs NHE) and lower than expected with 2-methoyxphenol (E° = 818 mV vs NHE), suggesting that an alkyl group *para* to the phenolic oxygen may be beneficial for substrate recognition. The activities of M298**C5a** and M298**C6a** with 4-methyl-2-methoxyphenol were similar to those with 4-ethyl-2-methoxyphenol (Supplemental Figure 6 and 7), indicating that the ethyl group is not specifically responsible for the improvements in activity. Instead, we posit that replacement of the axial methionine with bulky cycloalkyl-containing ncAAs is particularly beneficial for substrate recognition of *para*-alkylated phenols. Interestingly, such motifs are found in lignin, a prominent target for biomass valorization^26,27^.

Taken together, these results suggest that the ncAA mutations have a beneficial effect directly on the rate-limiting reaction step(s), which are often more challenging to improve compared to simple substrate recognition. For both high-redox and low-redox laccases (including *Sc*SLAC), it is typically reported that electron transfer from the substrate to the T1Cu center is rate-limiting^24,61,62^, supporting our hypothesis that the M298**C5a** and M298**C6a** mutations could improve catalysis through direct stabilization of the reduced T1Cu center.

### M298C5a and M298C6a prevent water accumulation near the T1Cu center

To understand the structure-function relationships that give rise to improvement with M298**C5a** and M298**C6a**, we turned to protein crystallography. We obtained crystallographic data of *Sc*SLAC M298**C6a** (Figure 4a-b, PDB: 9HU7). These structures were compared to previously reported structures for *Sc*SLAC M298, M298F, and M298L (Figure 4c-f, PDBs: 7BDN, 7B2K, and 7B4Y, respectively)^46^. Comparison of the *Sc*SLAC M298**C6a** structure to each of the other variants indicated that the overall structures are nearly identical (RMSDs: 0.218-0.234 Å). However, there is a striking difference at the T1Cu center. The orientation of the **C6a** side chain is directly over the T1Cu site. The resolution was not sufficient to distinguish between a chair, twist, or boat conformations of the cyclohexyl ring. Thus, the chair conformation was selected for refinement because it is thermodynamically more stable^63^. A space-filling view of the **C6a** side chain (Figure 4b) demonstrates the large volume of the side chain. The orientation of the **C6a** side chain is similar to that of the methionine side chain in wt *Sc*SLAC, which contains an elongated fourth coordinative bond between S_Met_-Cu (Figure 4c). In the case of M298L, the shorter side chain allows for the accumulation of an ordered water molecule near the copper center (∼4.5 Å). Although the side chain of M298F is larger than leucine, the planar phenyl ring of M298F is reoriented to flip outward, squeezing into a small pocket away from the T1Cu and towards the protein surface. This reorientation also allows for accumulation of an ordered water molecule near the T1Cu center (∼4.7 Å). Based on the packing of M298F in the reorientation pose, we propose that the **C5a** and **C6a** side chains prevent similar reorientations because of the bulky non-planar ring of these ncAAs (Supplemental Figure 8). Thus, the M298**C6a** mutation removes a coordinating ligand, increases the hydrophobicity around the T1Cu site, and also provides steric bulk to occlude water further increasing the hydrophobicity and potentially limiting the reorganization energy upon reduction from Cu^2+^ to Cu^1+^. Based on our findings, we propose that M298**C5a** and M298**C6a** exhibit improved activity through this multi-factorial stabilization of the Cu^1+^ state.

**Figure 4.**
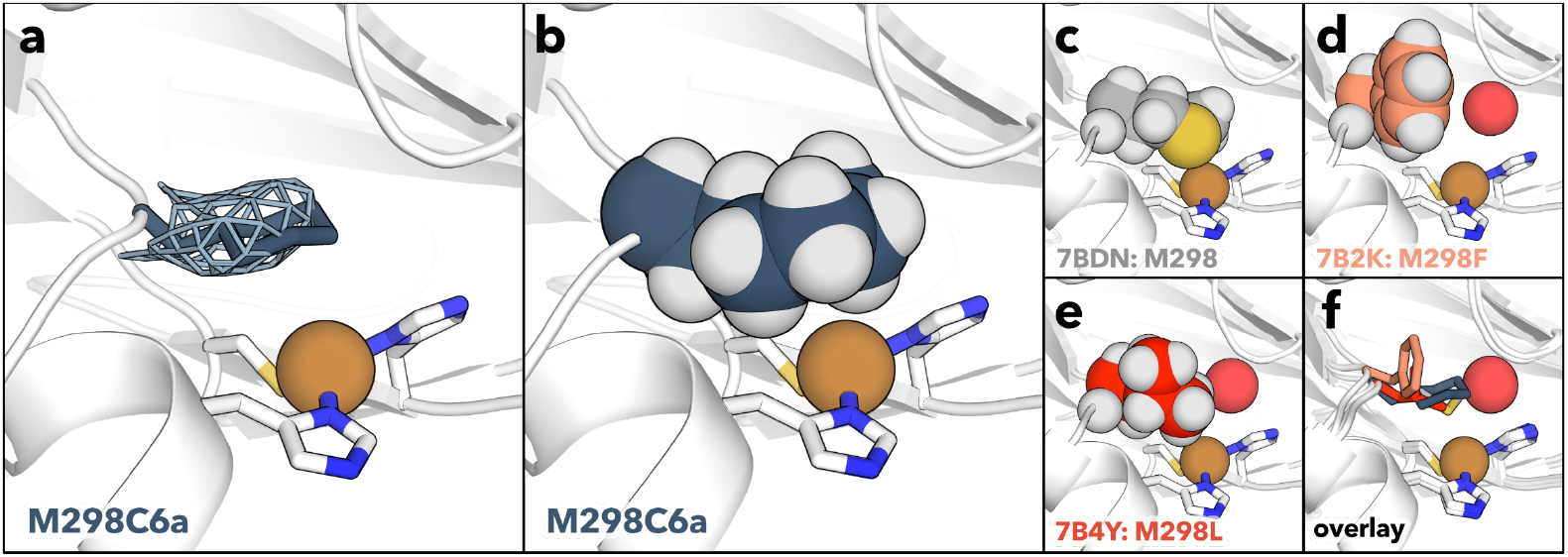
Structural comparison of *Sc*SLAC variants. The structure of *Sc*SLAC M298**C6a** was determined by x-ray crystallography (PDB: 9HU7). a) The 2Fo-Fc electron density map is rendered in mesh for the **C6a** side chain *(σ*: 1.0). Space-filling views for the different M298 variants are shown to demonstrate the differences between each mutation: M298**C6a** (b, 9HU7), wt *Sc*SLAC M298 (c, 7BDN), M298F (d, 7B2K), and M298L (e, 7B4Y). An overlay of the structures shows the relative orientations of the side chains and the potential steric clash between the order water molecules and the M298 and M298C6a (f). The copper ions are shown as orange spheres; the side chains of the axial amino acids are shown as spheres; and the ordered water molecules near the T1Cu copper in M298F and M298L are each shown as a single sphere for the oxygen atom. For all spheres, the relative scale is set to 70% of the van der Waals radii.

## Discussion

In this study, we developed a novel toolkit (PylRS^C6a^/PylT^m15^_CUA_) for the robust incorporation of a set of bulky, hydrophobic ncAAs: **C5a, C6a**, and **C7a**. In the absence of the target ncAAs, this system incorporates phenylalanine, but in the presence of the target ncAAs, the system shows high selectivity with no incorporation of canonical amino acids. For **C6a** incorporation, we obtained wt-like yields of protein using amber suppression for both sfGFP and *Sc*SLAC. Beyond the work presented herein, we expect that these new genetic code expansion tools will be valuable to explore and fine-tune the hydrophobicity of proteins in catalysis^5^ and binding interactions^64^. Additionally, because PylRS/PylT pairs can be typically shuttled between *E. coli* and eukaryotic cells^65^, the developed tools could be applied to tuning and studying the effects of hydrophobic tuning of a wide array of proteins. Moreover, owing to the use of **C6a** for tuning peptide properties, we anticipate that the ability to genetically incorporate this ncAA could have applications in peptide chemistry^66–68^.

With this robust method in hand, we leveraged the PylRS^C6a^/PylT^m15^_CUA_ pair to study and engineer the bacterial laccase *Sc*SLAC. A major drawback of bacterial laccases is their low activity, which is often linked to low redox potential of these enzymes. Mutations in the primary coordination-sphere have long been of interest for improving catalytic function. However, such modifications with traditional mutagenesis are highly detrimental to catalysis. Herein, semi-rational ncAA-based mutagenesis was used to improve the redox potential and catalytic activity of the bacterial laccase *Sc*SLAC through tuning the hydrophobic environment. We posit that this improvement in activity is derived from several factors: lowering the electron density at the copper center by removal of the coordinating methionine, increasing the hydrophobicity around the metal center, and providing steric bulk to physically prevent water from accumulating near the copper center. Notably, these improvements are derived without additional mutation in the secondary coordination sphere. However, the E°’_T1Cu_ has been suggested to be influenced by several factors including the geometry of the metal center, the metal-ligand interactions in the equatorial plane, the nature of the axial ligand, as well as the secondary coordination sphere hydrophobicity, hydrogen bonding, and electrostatics^69,19,70,3,71–73^. Thus, we expect that the *Sc*SLAC variants described herein will provide a valuable starting point for further improvements in both redox potential and enzymatic activity through semi-rational engineering around the T1Cu site^43^, computationally-guided engineering^74,75^, and directed evolution^76^. Additional mutants with improved activity could enable unprecedented applications for these enzymes.

Mutagenesis with ncAAs has shown continued promise for enzymatic characterization and enzyme engineering. Most ncAA-based enzyme engineering focuses on improved non-native activity, often through improving substrate recognition, improving enantioselectivity, or installing an abiotic catalytic entity. Here, we demonstrate that ncAA-based enzyme engineering can be used to improve near-native catalysis and directly improve the rate-limiting chemical step(s) through semi-rational design using fundamental chemical principles. The ability to use ncAA-based enzyme engineering to overcome a multidecade effort towards engineering bacterial laccases demonstrates the power of this technique. Our findings suggest that with the continued discovery of robust translation systems for ncAA incorporation, ncAA-based enzyme engineering will become not just more viable alternative, but also highly valuable tool for enzyme engineering.

## Methods

### Mass spectrometry methods

During aaRS/tRNA screening, initial high-throughput liquid chromatography-mass spectrometry (LC-MS) of sfGFP was conducted using an Agilent single quadrupole mass spectrometer (ESI), coupled to a 1290 Infinity II LC system.

Verification of all proteins discussed in the text was additionally carried out by high-resolution LC-MS carried out at the Functional Genomics Center Zürich (FGCZ). Briefly, samples were resolved on an ACQUITY UPLC@BioResolve-RP-mAb (2.7 μm, 2.1 mm x 150 mm, 450 Å) column. The analysis was performed on a Synapt G2-Si mass spectrometer directly coupled to the UPLC station. Mass spectra were acquired in the positive-ion mode by scanning the m/z range from 400 to 5000 Da with a scan duration of 1 s and an interscan delay of 0.1 s. The spray voltage was set to 3 kV, the cone voltage to 50 V, and the source temperature to 100 °C. The data were recorded with the MassLynx 4.2 Software. The recorded m/z data of single peaks were deconvoluted into mass spectra by applying the maximum entropy algorithm MaxEnt1 (MaxLynx) with a resolution of the output mass 0.5 Da/channel and Uniform Gaussian Damage Model at the half height of 0.7 Da.

LC-MS/MS was carried out by the FGCZ. Purified protein (∼ 1 mg/mL) was digested in solution by mixing 5 μL of sample with 40 μL digestion buffer (10 mM Tris, 2 mM CaCl_2_, pH 8.2). Protein was reduced and alkylated by 0.9 μL 100 mM TCEP + 1.4 μL 100mM ClAA. 2 μL trypsin (100 ng/μL in 10 mM HCl) were added and microwave assisted digestion was carried out at 60 °C for 30 min. The samples were dried and dissolved in 20 μL ddH_2_O + 0.1% formic acid. Samples were analyzed on M class UPLC coupled to a Q-Exactive mass spectrometer (Thermo). Data were searched against sfGFP sequence sequences by Byonic 5.2 (Proteinmetrics, USA).

Small-molecule LC-MS analysis and UPLC analysis were both conducted with an Agilent 1290 Infinity II LC system coupled to a single quadrupole mass spectrometer (ESI).

### General instrumentation and materials

UV-VIS were measured either on a Multiskan Sky (Thermo Scientific) or a Cary 60 UV-Vis (Agilent), except for kinetics measurements, which were measured on a Tecan Infinite M Nano+. ^1^H-NMR spectra were recorded in CDCl_3_ or D_2_O on a Bruker AV-AV-400 (400 MHz), chemical shift d in ppm relative to solvent signals (d = 7.26 ppm for CDCl_3_, 4.79 ppm D_2_O), coupling constants J are given in Hz. ^13^C-NMR spectra were recorded in D_2_O on a Bruker AV-AV-400 (400 MHz). All purchased chemicals were used without further purification. Automated flash chromatography was conducted on a Biotage Isolera instrument. All sequencing was conducted by Microsynth (Balgach, CH).

### aaRS Library construction

Libraries were created using modified Golden Gate cloning. A pSL plasmid encoding the corresponding synthetase was amplified using inverse PCR with primers carrying a BsaI restriction site and an additional 6 bp overhang at the 5’ end. PCR products were purified using an NEB Monarch PCR cleanup kit. The purified PCR products were digested with DpnI and BsaI in 1x rCutSmart. Digests were carried out at 37 °C overnight. Digested products were purified using an NEB Monarch PCR cleanup kit. DNA ligation reactions contained T4-ligase and 1x T4-ligase-buffer. Ligation was carried out at 16 °C overnight. Ligation products were purified using an NEB Monarch PCR cleanup kit. Electrocompetent NEB10β *E. coli* (100 μL) were transformed with ligation product by electroporation in a cuvette with 2 mm gap on an Eppendorf Eporator with 250 Ω and 2500 V with pulse times of ∼5 ms. The cells were recovered in SOC media for 1 h at 37 °C with shaking at 220 rpm. The number of transformants was estimated by dilution series plating of LB-agar with 50 μg/mL kanamycin. The recovered cells were transferred to 10 mL LB media with 50 μg/mL kanamycin and grown for 5 h. The cells were harvested by centrifugation, and the DNA was isolated using an NEB Monarch Plasmid Miniprep Kit. The library quality was confirmed by Sanger sequencing of 2-3 individual clones.

### aaRS Library selections

For positive selections, electrocompetent NEB10β cells (100 μL) carrying a pDPS2-*Mm*PylT_CUA_ plasmid were transformed with 250 ng of a given library by electroporation in a cuvette with 2 mm gap on an Eppendorf Eporator at 250 Ω and 2500 V with pulse times of ∼5 ms. The cells were recovered in SOC media for 1 h at 37 °C with shaking and transferred to 10 mL LB media with 50 μg/mL kanamycin, 10 μg/mL tetracycline, and 6 mM **C6a**. The cells were grown for 1-2 h, harvested by centrifugation, resuspended in 250 μL LB media and plated on LB-agar with 0.4% arabinose, 50 μg/mL kanamycin, 10 μg/mL tetracycline, 100 μg/mL chloramphenicol, 0.4% arabinose, and 6 mM **C6a**. Plates were incubated for 24 h at 37 °C. If additional selection rounds were carried out, cells were collected by washing the plate with 10 mL LB media and harvesting the cells by centrifugation. Plasmid DNA was isolated using an NEB Monarch Plasmid Miniprep Kit. DNA was digested with AgeI for 18h at 37 °C to remove the selection plasmid and purified using an NEB Monarch PCR cleanup kit. If additional selection rounds were not carried out, we proceeded as though the step was the final positive selection round described further below.

For negative selections, electrocompetent NEB10β cells (100 μL) carrying a pBARN-*Mm*PylT_CUA_ plasmid were transformed with 50 ng of library DNA by electroporation in a cuvette with 2 mm gap on an Eppendorf Eporator at 250 Ω and 2500 V with pulse times of ∼5 ms. The cells were recovered in SOC media for 1 h at 37 °C with shaking and transferred to 10 mL LB media with 50 μg/mL kanamycin and 35 μg/mL chloramphenicol. The cells were grown in the presence of the antibiotics for 1-2 h. The cells were harvested by centrifugation, resuspended in 250 μL LB media and plated on LB-agar with 0.4% arabinose, 50 μg/mL kanamycin, and 35 μg/mL chloramphenicol. Plates were incubated for 24 h at 37 °C. Cells were collected by washing the plate with 10 mL LB media; cells were harvested by centrifugation; and plasmid DNA was purified using an NEB Monarch Plasmid Miniprep Kit. DNA was digested with AgeI in 1x rCutsmart. Digest were carried out at 37 °C overnight to the remove selection plasmid and purified using an NEB Monarch PCR cleanup kit.

For screening after the final round of positive selection, individual colonies were used to inoculate 96-well plates with 250 μL 2xYT media with 50 μg/mL kanamycin, 10 μg/mL tetracycline. Cultures were grown for 24 h at 37 °C with shaking at 400 rpm. Subsequently, 25 μL of each culture were used to inoculate 96-well plates with 250 μL 2xYT media with 50 μg/mL kanamycin, 10 μg/mL tetracycline, 0.4% arabinose, and ± 6 mM **C6a**. Cultures were grown for 24 h at 37 °C with shaking at 400 rpm. 100 μL of each culture were transferred to a 96-well clear well plate. Fluorescence (excitation at 480 nm and emission at 510 nm) and absorbance at 600 nm were measured. DNA from cultures with high Fluorescence/OD_600_ ratios was isolated using an NEB Monarch Plasmid Miniprep Kit and analyzed by Sanger sequencing to identify mutations in the PylRS gene.

### sfGFP amber suppression assay

NEB10β *E. coli* were co-transformed with pBAD-sfGFP150_TAG_ (modified from Addgene #85483)^77^ and the corresponding aaRS/tRNA expression plasmid by heat shock (42 °C, 30 s), recovered in SOC for 1 h at 37 °C and plated on LB-agar with 50 μg/mL kanamycin and 10 μg/mL tetracycline. Plates were incubated for 24 h at 37 °C. Individual colonies were used to inoculate 250 μL autoinduction media with 100 μg/mL kanamycin, 20 μg/mL tetracycline, and ncAA. For each ncAA the concentrations were as follows: 6 mM **C6a** and 12 mM dl-ncAA for **C5a** and **C7a**. wt sfGFP was analogously expressed from pBAD-sfGFP by omission of 10 μg/mL tetracycline addition for the culturing of cells. 100 μL of each culture were transferred to a 96-well clear well plate. Fluorescence (excitation at 480 nm and emission at 510 nm) and absorbance at 600 nm were measured.

### Analysis of ncAA incorporation in sfGFP

NEB10β *E. coli* were co-transformed with pBAD-sfGFP150_TAG_ and the corresponding aaRS/tRNA expression plasmid by heat shock (42 °C, 30 s), recovered in SOC for 1 h at 37 °C and plated on LB-agar with 50 μg/mL kanamycin and 10 μg/mL tetracycline. Plates were incubated for 24 h at 37 °C. Individual colonies were used to inoculate 5 mL autoinduction media with 100 μg/mL kanamycin, 20 μg/mL tetracycline, and either 12 mM DL-**C5a**, 6 mM **C6a**, or 12 mM DL-**C7a**. The cultures were grown for 24 h at 37 °C with shaking at 240 rpm. The cells were harvest by centrifugation for 10 min at 4 °C and 4200 g; the supernatant was decanted; and the cell pellets were stored at -80 °C. To isolate the sfGFP, the cells were thawed at room temperature and resuspended in lysis buffer (500 μL, 20 mM Tris, 300 mM NaCl, pH 7.2 at 4 °C, 0.2% *n*-octyl *beta*-D-thioglucopyranoside, 4 mg/mL Lysozyme). The lysis was conducted at 22 °C for 4 h. The sfGFP was isolated by purification with Ni-NTA resin (HisPur from Thermo Scientific) according to the manufacture’s instructions. The purified sfGFP was analyzed by LC-MS as indicated in the Mass Spectrometry section.

### *Sc*SLAC production

The N-terminal signaling peptide (amino acids 2-30) of *Sc*SLAC is highly prone to proteolysis.^78^ In laccases, this region is not necessary for catalysis. In many laccase studies—including *Sc*SLAC studies, the N-terminal region is truncated^39,76,78,79^. Thus, to reduce complexity and ensure that LC-MS could be used to identify ncAA incorporation, we designed an *Sc*SLAC construct without the N-terminal signaling peptide and with a C-terminal His6 tag for purification. Both constructs yielded similar catalytic parameters with ABTS, which were similar to previous literature reports with similar conditions^43^. Thus, we continued with the N-terminally truncated *Sc*SLAC-His6. The sequence is provided in the DNA and protein sequence section.

To produce *Sc*SLAC, the corresponding pET28-SLAC-M298X (where X is M, L, or F) plasmid was transformed into B-95ΔAΔfabR cells (Addgene #197934)^53^. Precultures were grown at 37 °C overnight in LB with 50 μg/mL kanamycin from single colonies or glycerol stocks. The cells were diluted 100-fold into 25-50 mL autoinduction media with 50 μg/mL kanamycin in a baffled flask and shaken at 37 °C for 3 h. CuCl_2_ was added to a final concentration of 2 mM, and the cells were shaken (160 rpm) at 25 °C for 28 h. The cells were collected by centrifugation (10 min, 4 °C, 4200 g), the supernatant was decanted, and the cells were frozen at -80 °C.

### *Sc*SLAC production with ncAAs

B-95ΔAΔfabR cells (Addgene #197934)^53^ were transformed with pGS1T-PylRS^C6a^-PylT^m15^_CUA_ and the desired pET28-SLAC-M298_TAG_ plasmid. Precultures were grown at 37 °C overnight in LB with 50 μg/mL kanamycin and 10 μg/mL tetracycline from single colonies or glycerol stocks. The cells were diluted 100-fold into 50-100 mL autoinduction media with 50 μg/mL kanamycin, 10 μg/mL tetracycline, and either 20 mM DL-**C5a**, 8 mM **C6a** or 12 mM DL-**C7a** in a baffled flask and shaken at 37 °C for 8-14 h. CuCl_2_ was added to a final concentration of 2 mM, and the cells were shaken (160 rpm) at 25 °C for 24-28 h. The cells were collected by centrifugation (10 min, 4 °C, 4200 g); the supernatant was decanted; and the cells were frozen at -80 °C.

### *Sc*SLAC Purification

Cells were thawed on ice and lysed with lysis buffer (20 mL/g cell pellet, 20 mM Tris, 500 mM NaCl, pH 8.0 at 4 °C, 0.25% *n*-octyl *beta*-D-thioglucopyranoside, 2 mg/mL Lysozyme, 0.25 μL/mL DNase, 2 mM MgCl_2_) at 4 °C overnight, turning end-over-end. CuCl_2_ was added to a final concentration of 2 mM, and the lysate was turned end-over-end at 4 °C for 3 h. The lysate was pelleted by centrifugation (18000 g, 4 °C, 1 h), and the supernatant was decanted. Ni-NTA beads (1 mL/g cell pellet) were added to the supernatant, and the suspension was turned end-over-end at 4°C for 30-45 min. The Ni-NTA beads were collected by centrifugation (100 g, 4 °C, 5 min); the supernatant was decanted; and the beads were loaded onto a gravity flow column. The columns were washed four times with 4 mL of washing buffer (20 mM HEPES, 300 mM NaCl, 20 mM imidazole, pH 7.0), and the protein was collected with elution buffer (20 mM HEPES, 300 mM NaCl, 500 mM imidazole, pH 7.0). Fractions with protein were combined, and the protein was buffer exchanged to storage buffer (20 mM HEPES, 300 mM NaCl, pH 7.0) using a Zeba™ Spin Desalting Column (2 mL or 5 mL, 7 kDa MWCO). Fractions with protein were pooled, flash frozen in liquid nitrogen and stored at -80 °C.

### Crystallization and structure determination

*Sc*SLAC M298**C6a** was concentrated to 18-20 mg/mL using Amicon Ultra-4 centrifugal filter tubes (10 kDa molecular-weight cutoff, Merck Millipore) in storage buffer (20 mM HEPES, 300 mM NaCl, pH 7.0). Crystals of ScSLAC M298**C6a** were obtained at 20 °C within 4 weeks by sitting-drop vapor diffusion with *Sc*SLAC M298**C6a** (150 nL) and a precipitant of 18.2% Jeffamine-M600 and 17.5% DMSO (100 nL precipitant). The crystals were mounted onto MicroMesh (400/10 μm, MiTiGen) and flash-cooled in liquid nitrogen after the addition of 25% glycerol as cryoprotectant.

Data were collected at ESRF MASSIF-3/ID30-A3 Beamline at 0.9677Å. Multiple datasets of one crystal were collected due to weak diffraction. The data was processed using the Autoprocess pipeline^80–85^. The data could be merged and scaled to 3.49 Å and were solved by molecular replacement using the Alphafold3^86^ model of the protein. The crystal contained one molecule per asymmetric unit with a high solvent content of >85% that may have contributed to the weak diffraction but may have aided bulk solvent correction during crystallographic refinement. The initial model was refined and rebuilt iteratively in Coot^87^ and phenix.refine^88^. *Sc*SALC M298**C6a** could be refined to an R_work_/R_free_ 0.1874/0.2305 and a Molprobity score of 1.22. The map was of exceptional quality (Supplementary Figure 8a). The PDB model has been deposited in the PDB as 9HU7. In the PDB deposition, **C6a** was termed ALC based on previous PDB entries. Crystallographic details are provided in Supplementary Table 2.

### Michaelis-Menten kinetics

The reactions were carried out in 20 mM Britton-Robinson (BR) buffer (20 mM acetic acid, 20 mM phosphoric acid, 20 mM boronic acid, pH adjusted with aq. NaOH) at pH 4.0. To prepare the reactions, 2X solutions of reaction components were prepared as follows: 2X BR buffer, 2 mM CuCl_2_, and varying ABTS concentrations (0.2, 0.5, 1, 2, 4, 6, 8, or 10 mM). Separately, the desired enzyme was diluted to 400 nM in water for wt, **C5a**, and **C6a** and to 2000 nM for M298L and M298F. To initiate the reaction, 50 μL of the enzyme solution were added to 50 μL of the 2X reaction component solution. As a control, samples without enzyme were prepared with 50 μL of water instead of the 50 μL of dilute enzyme. Immediately after mixing by pipetting up and down, the absorption change at 420 nm (ε_420_ = 36000 M^-1^cm^-1^) was measured every 9-10 s. The reaction was carried out at 25 °C. For data processing, for each ABTS concentration, the time trace corresponding to the negative control (without enzyme) was subtracted from each time trace containing protein. The kinetic parameters were determined by Michaelis-Menten analysis.

### Redox potential determination

Redox potentials were determined by redox titration using K_3_[Fe(CN)_6_] as redox dye and sodium dithionite as reductant. The titrations were performed in a glovebox under nitrogen atmosphere. The following components were mixed together in a total volume of 250 μL: 110-285 μM protein, 500 μM K_3_[Fe(CN)_6_], 4 μM methyl viologen in 20 mM HEPES, 300 mM NaCl, pH 7.0. The absorbance spectra were measured from 300-900 nm after every addition of 0.5-2 μL sodium dithionite (5 mM) and an equilibration time of 20 min. A baseline correction was made based on absorbance at 900 nm. The amount of oxidized protein and dye was determined by spectral deconvolution by fitting linear combinations of the spectra of oxidized protein and dye from 380 – 650 nm. The reduced species did not absorb within the wavelength range of interest. The percentage of oxidized and reduced dye was used to determine the redox potential of the system using the Nernst equation as follows.

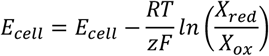

At equilibrium, the two half reactions can be described as follows.

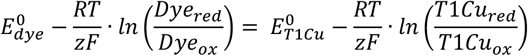

A redox potential of 436 mV vs NHE for K_3_[Fe(CN)_6_] was used^43^, and the conditions for RT/zF were R = 8.314 J•K^-1^•mol^-1^, T = 293 K, F = 96485 C•mol^-1^, and z = 1.

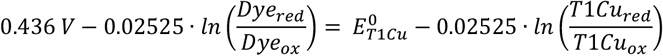

At each titration point, the deconvolution data provides the concentration of the reduced and oxidized dye and the concentration of the reduced and oxidized T1Cu center. With this information, we can treat the equation as a linear function where the left side of the equation is y, **E°’**_**T1Cu**_ **is the y-intercept (b)**, RT/zF is the slope (m), and ln(T1Cu_red_/T1Cu_ox_) is x.

### Total turnover number measurements

The reactions were carried out in 20 mM Britton-Robinson (BR) buffer (20 mM acetic acid, 20 mM phosphoric acid, 20 mM boronic acid, pH adjusted with aq. NaOH) at pH 7.0. The following reaction components were mixed together in a reaction volume of 100 μL: 970 μM phenolic substrate, 970 μM CuCl_2_, 11.1-1388 nM enzyme^⫩^ in 20 mM BR buffer. Reactions omitting enzyme were prepared as negative controls. The reaction was carried out at 25 °C for 24 h while shaking at 500 rpm in an Accutherm microtube shaking incubator. The reaction was quenched by the addition of 10 μL aq. HCl (2 M). The reaction mixture was clarified by centrifuged (5 min, 14000 g), and 50 μL supernatant was diluted in 100 μL water. The mixture was analyzed by UPLC. The UV-VIS peak (220 nm) corresponding to the desired phenolic substrate was integrated, and the remaining, unconverted substrate concentration was determined by comparison to a standard calibration curve. The conversion was then calculated as the concentration of substrate in the negative control minus the concentration of unconverted substrate. This final enzymatic conversion was then divided by enzyme concentration to determine the total turnover number (TTN).

^⫩^For substrates with higher redox potentials, the enzyme concentrations were increased to ensure that the product concentration was well within the detection limit of our assay.

## Acknowledgments

The authors thank the University of Zurich for funding, the Functional Genomics Center Zurich (FGCZ) for assistance with high-resolution LC-MS and LC-MS/MS data collection and analysis, the European Synchrotron Radiation Facility (ESRF) for provision of synchrotron radiation facilities (MX2644), the staff of the ESRF and EMBL Grenoble for assistance and support in using MASSIF-3/ID30-A3 (MX2644), the UZH Protein crystallization center for assistance with crystallization, and the Protein Production and Structure Core Facility (EPFL) for assistance with data collection and analysis for crystallography. The authors thank the laboratory of Eva Freisinger for access to the atomic absorption spectrometer and the anaerobic chamber used for redox titrations. SF also acknowledges the UZH CanDoc Award for funding (grant no. FK-22-086).

## Author contributions

ADL conceived of and supervised the project. ADL, SF, and ANP contributed to design of experiments. SF synthetized **C5a** and **C7a**. ANP engineered and validated the aaRS/tRNA pairs for incorporation of **C5a, C6a**, and **C7a** into sfGFP. SF and ANP expressed the enzyme variants. SF characterized the enzyme variants and prepared crystals for M298**C6a**. KL performed structure refinement for M298**C6a**. All authors contributed to writing and editing the final paper.

