## Supplementary Information for "Hydrophobic tuning with non-canonical amino acids in a copper metalloenzyme"

**Table of Contents**

| | | |
|---|---|---|
| I. | Supplementary data figures | 2 |
| II. | Synthesis | 10 |
| III. | Kinetic parameters | 17 |
| IV. | Crystallography | 18 |
| V. | References | 19 |

### I. Supplementary data figures

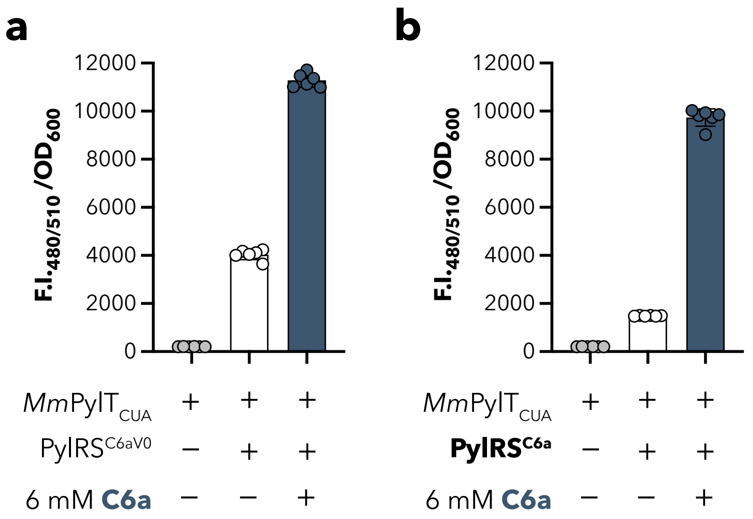

**Supplementary Figure 1 | Validation of PylRS variants for incorporation of C6a**. Suppression of sfGFP150$_{TAG}$ in NEB10β in the presence or absence of PylRS$^{C6a_v0}$ and **C6a** (a) and in the presence or absence of PylRS$^{C6a}$ and **C6a** (b). Data shows Fluorescence (excitation at 480 nm and emission at 510 nm) normalized to the absorbance at 600 nm and represents the mean and standard deviation of 6 biological replicates.

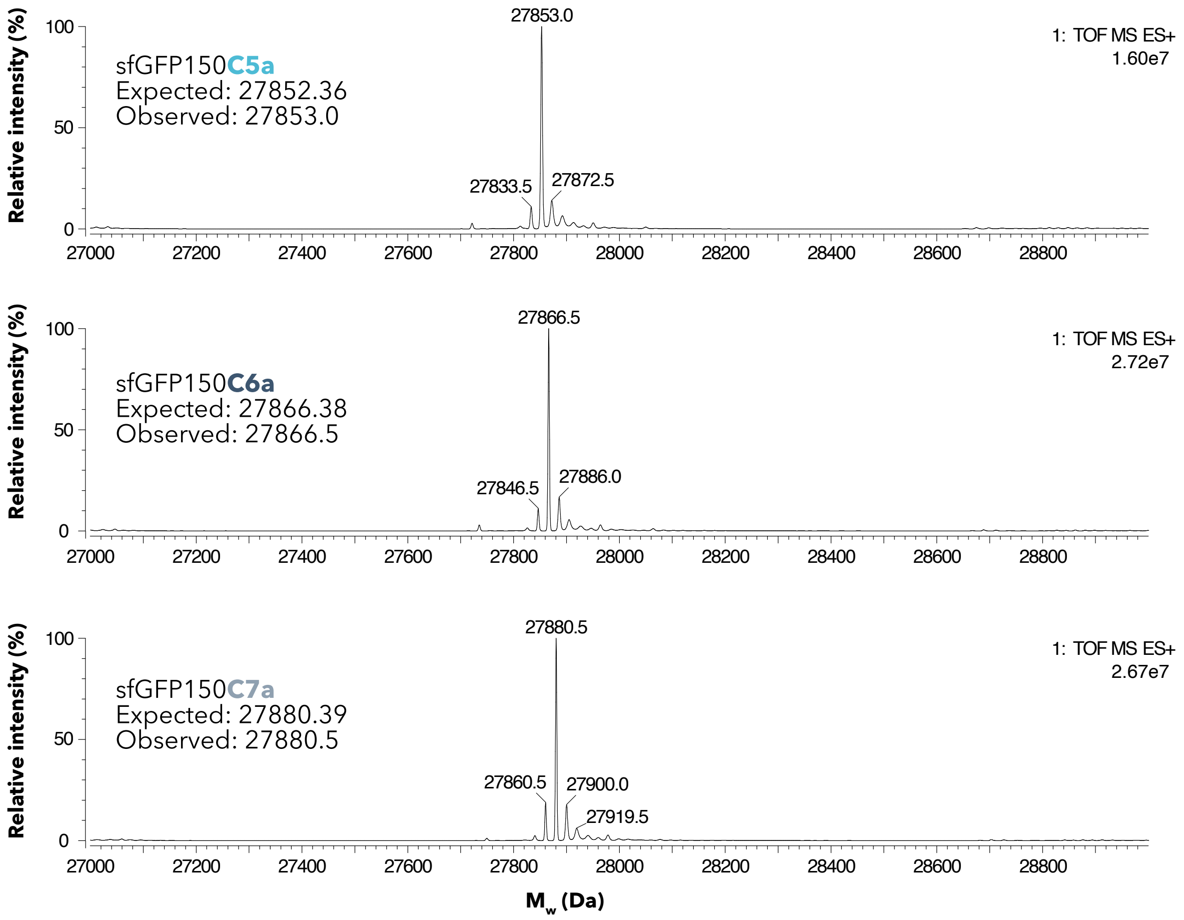

**Supplementary Figure 2 | Full deconvolution spectra for LC-MS analysis of sfGFP150$_{TAG}$ containing C5a, C6a, or C7a.** Spectra were collected for sfGFP150$_{TAG}$ expressed with pGS1T-PylRS$^{C6a}$-PylT$^{m15}$$_{CUA}$ in the presence of 12 mM of DL-**C5a** (top), 6 mM of **C6a** (middle), and 12 mM of DL-**C7a** (bottom).

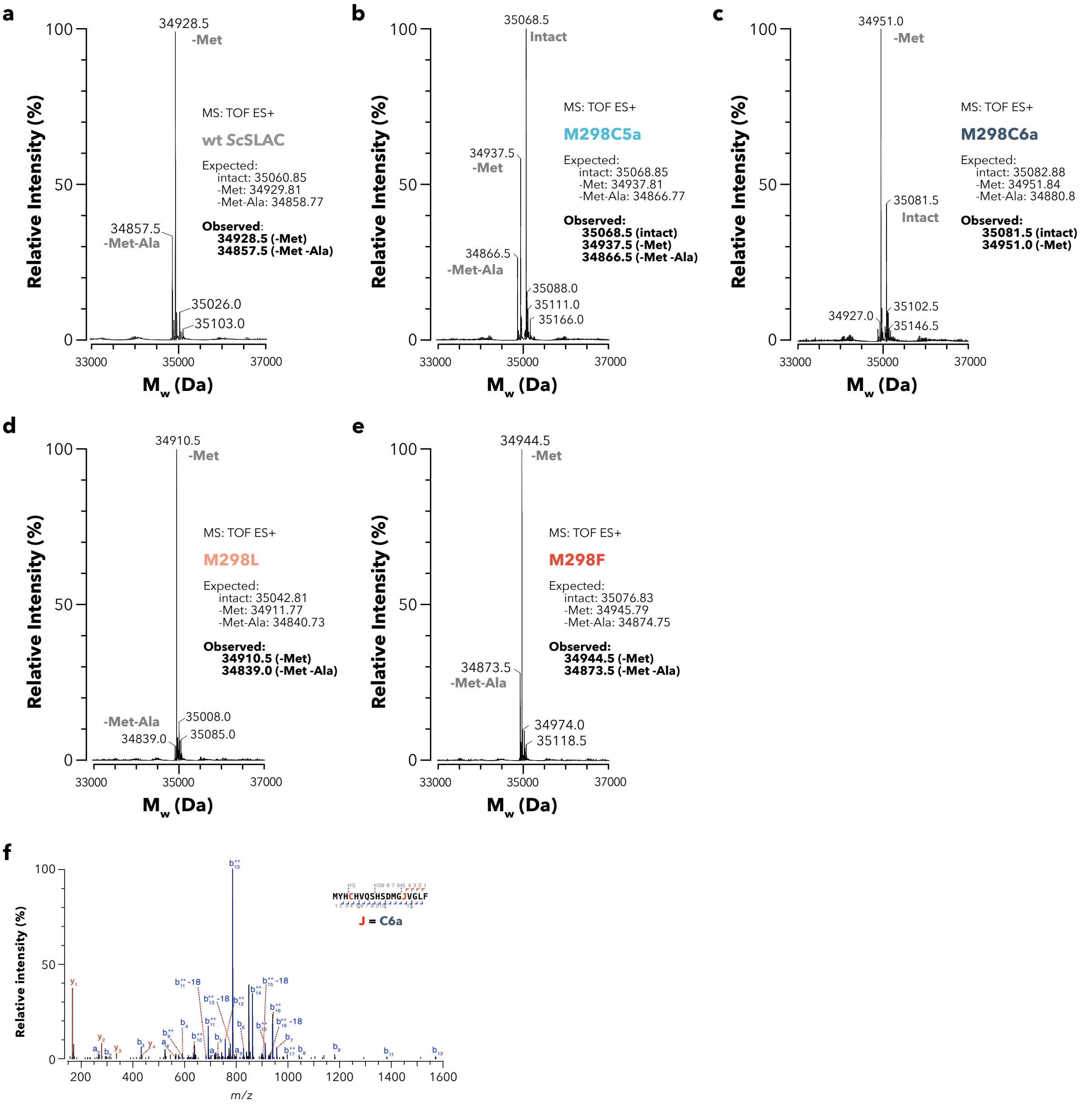

**Supplementary Figure 3 | LC-MS and LC-MS/MS analysis of *Sc*SLAC-M298-variants.** Mass spectra from LC-MS analysis of *Sc*SLAC variants: M298 (a), M298**C5a** (b), M298**C6a** (c), M298F (d), and M298L (e). Often truncation of the N-terminal Met and MetAla is observed. f) A representative LC-MS/MS spectrum from a tryptic digest of M298**C6a**. Over 100 peptides containing **C6a** were observed, but no peptides for canonical amino acid incorporation were observed.

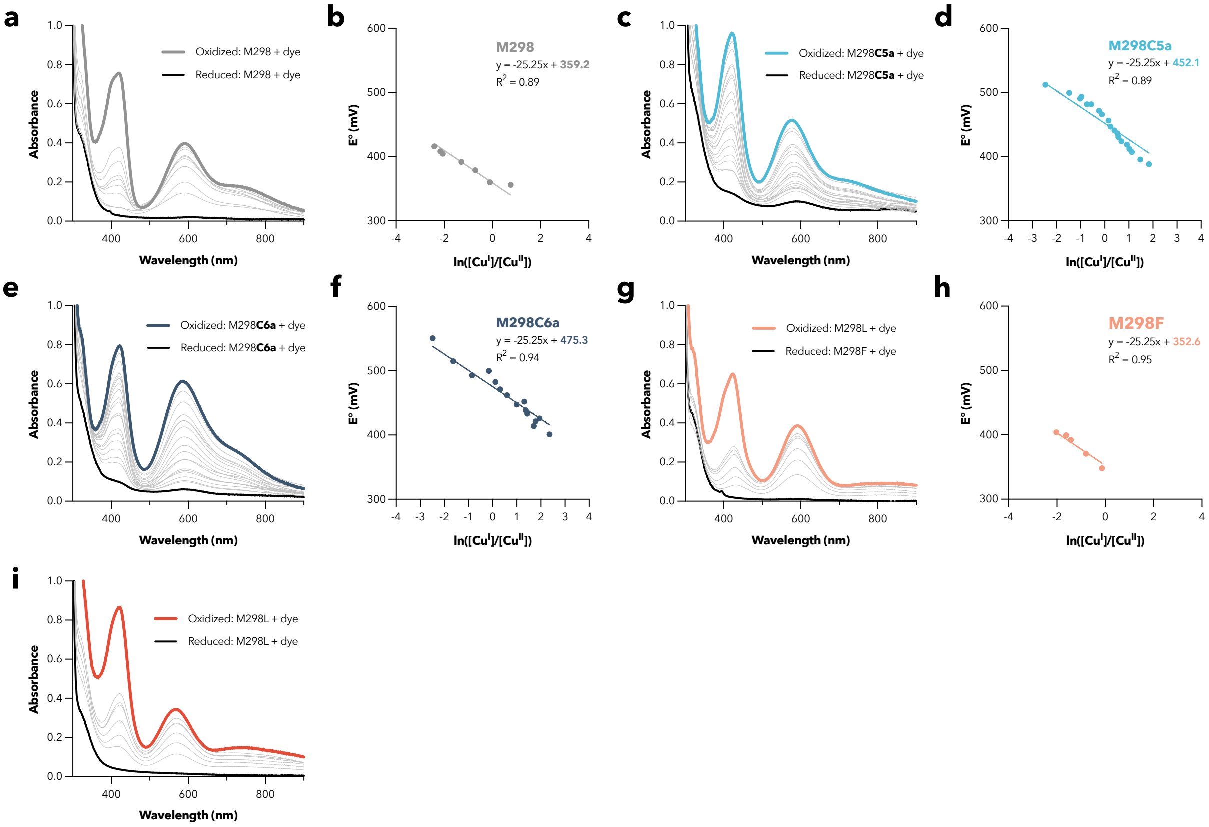

**Supplementary Figure 4 | Redox titrations of the different *Sc*SLAC variants: M298, M298C5a, M298C6a, M298F, and M298L.** For each variant, the UV-VIS spectra for each titration point is shown (a, c, e, g, and i). With these spectra, a deconvolution was conducted to calculate the ratio of reduced and oxidized dye and the ratio of reduced and oxidized T1Cu site. The potential of the cell ($E^0$) was plotted as a function of the ln($Cu^{1+}/Cu^{2+}$), insets b, d, f, and h. The redox potential of M298L was very low, and so the dye was immediately reduced before the protein. Thus, it was not possible to determine the ratios of reduced and oxidized copper for this titration. A linear fit was applied according to Nernst equation: $E^0_{dye} - \frac{RT}{zF} \cdot ln\left(\frac{Dye_{red}}{Dye_{ox}}\right) = E^0_{T1Cu} - \frac{RT}{zF} \cdot ln\left(\frac{T1Cu_{red}}{T1Cu_{ox}}\right)$. The slope was fixed at a value of 25.25 corresponding to 1000RT/nF. The estimated redox potential of the T1Cu site is then given as the y-intercept (see Methods for a mathematical explanation). The factor of 1000 in the slope analysis is to account for the change from V to mV.

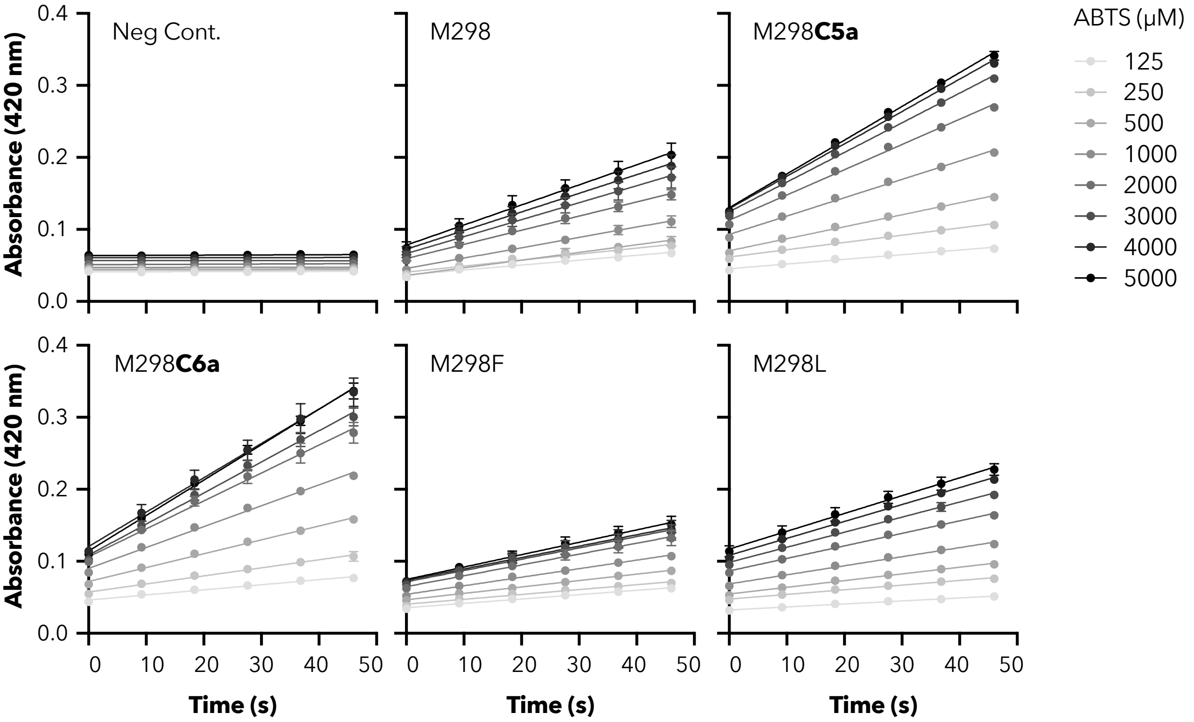

**Supplementary Figure 5 | Initial velocity data for kinetics analysis with ABTS.** The reactions were conducted in 20 mM BR buffer at pH 4.0, 1 mM $CuCl_2$, and varying ABTS concentrations. For M298, M298**C5a**, and M298**C6a**, the enzyme concentrations were 200 nM. For M298L and M298F, the enzyme concentrations were 1000 nM. For each enzyme, at least three biological replicates were collected. Three preparative replicates were conducted for the negative control, which omits enzyme but contains 1 mM $CuCl_2$. The error bars represent the standard deviation.

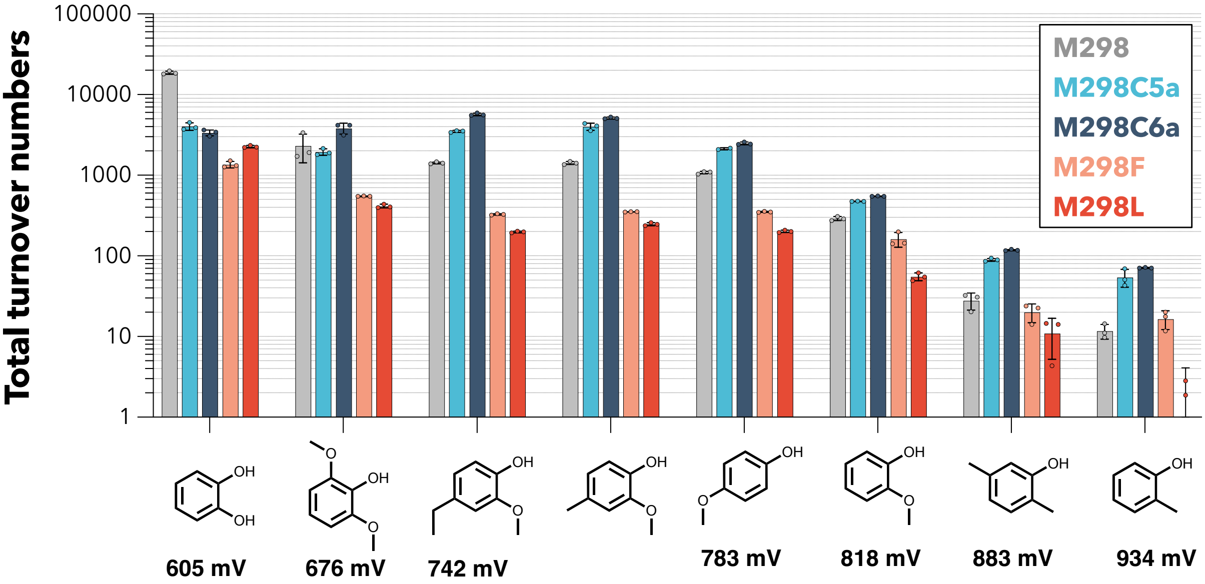

**Supplementary Figure 6 | Total turnover numbers with phenolic substrates.** Total turnover numbers of *Sc*SLAC M298, M298**C5a**, M298**C6a**, M298F, and M298L with different phenolic substrates with their corresponding redox potential vs NHE.[60] For internal consistency, the reported redox potentials were derived from a single publication, which used a standardized method for all measurements.[60] The redox potential of 4-methyl-2-methoxyphenol was not reported by this method. Thus, the redox potential is omitted for this species. However, we posit that it is near that of 4-ethyl-2-methoxyphenol based on other reports. Reactions were performed in biological triplicates and the error bars represent the standard deviation.

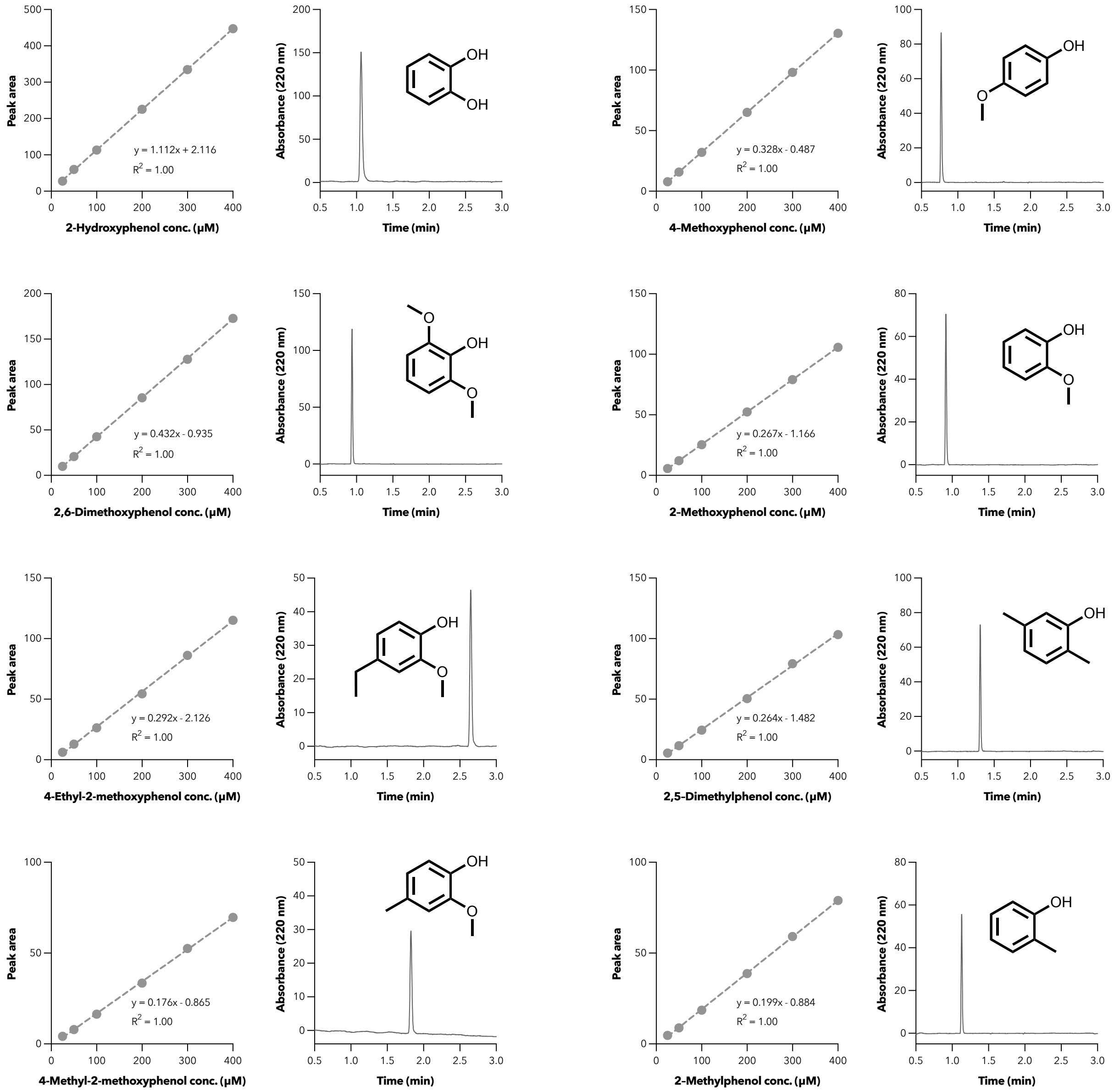

**Supplementary Figure 7 | Calibration curves and representative LC-MS trace for each phenolic substrate.** Calibration curves were prepared by integrating the peak area for standards at varying concentrations. All representative LC-MS data are for 300 µM phenolic substrate. All TTN experiments yielded substrate concentrations within the range of the calibration curves.

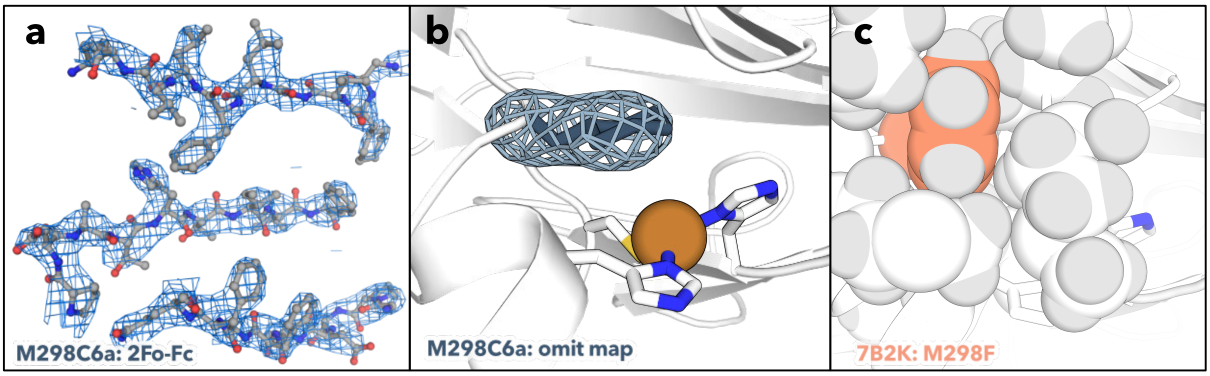

**Supplementary Figure 8 | Crystallographic comparisons of ScSLAC M298C6a and M298F.** a) The 2Fo-Fc map for *Sc*SLAC M298**C6a** (PDB: 9HU7) illustrating the map quality, contoured at $\sigma$ = 1.0. b) The omit map for residue M298**C6a** indicating contoured at σ = 8.0, indicating that the unassigned density fits well to **C6a**. c) A space filling view of the M298F packing, which suggests that the area occupied by the phenyl ring is too narrow for the bulkier cyclohexyl side chain.

### II. Synthesis

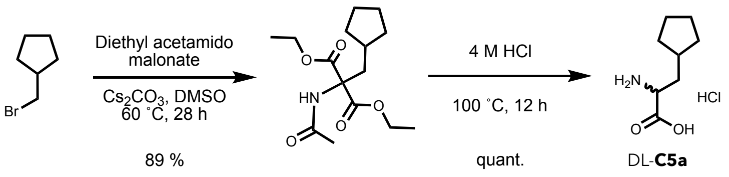

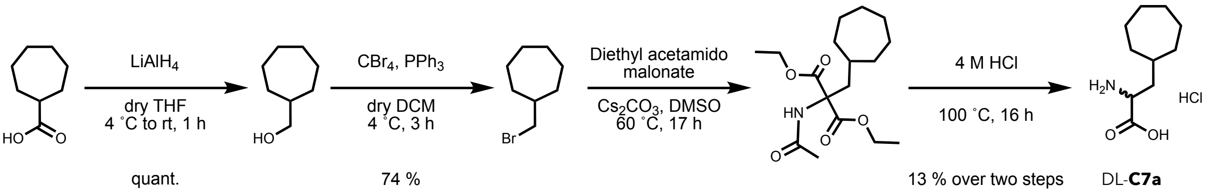

#### Diethyl 2-acetamido-2-(cyclopentylmethyl)malonate[89]

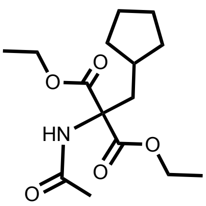

Diethyl acetamido malonate (769 mg, 3.54 mmol, 1.0 eq) was dissolved in DMSO (4 mL) under a nitrogen atmosphere. $Cs_2CO_3$ (1.73 g, 5.31 mmol, 1.5 eq.) was added, followed by the addition of cyclopentylmethylbromide (500 µL, 3.89 mmol, 1.1 eq.). The suspension was stirred at 60 °C for 28 h. The suspension was allowed to reach room temperature, and ice-cold water (50 mL) was added. The mixture was extracted with EtOAc (2 x 50 mL), and the combined organic phases were washed with brine (80 mL). The organic phase was dried over $Na_2SO_4$, and the supernatant was collected by filtration. The solvent was removed under reduced pressure to yield diethyl 2-acetamido-2-(cyclopentylmethyl)malonate (943 mg, 3.15 mmol, 89%) as a white solid.

$^1$H NMR (400 MHz, $CDCl_3$) δ 6.81 (s, 1H), 4.27 – 4.18 (m, 4H), 2.44 (d, $^3J_{HH}$ = 6.6 Hz, 2H), 2.03 (s, 3H), 1.76 – 1.38 (m, 7H), 1.25 (t, $^3J_{HH}$ = 7.1 Hz, 6H), 1.12 – 0.99 (m, 2H).

#### Cyclopentylalanine hydrochloride (DL-C5a)

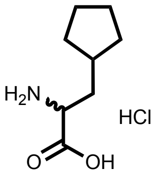

Diethyl 2-acetamido-2-(cyclopentylmethyl)malonate (943 mg, 3.15 mmol, 1.0 eq.) was suspended in aq. HCl (4 M, 20 mL) and stirred at 100 °C for 12 h. The solvent was removed under reduced pressure. The resulting white solid was dissolved in water, and the solution was lyophilized to yield DL-cyclopentylalanine hydrochloride (610 mg, 3.15 mmol, quant.) as a white powder.

$^1$H NMR (400 MHz, $D_2O$) δ 4.03 (t, $^3J_{HH}$ = 6.6 Hz, 1H), 2.08 – 1.78 (m, 5H), 1.73 – 1.50 (m, 4H), 1.26 – 1.11 (m, 2H).
$^{13}$C NMR (101 MHz, $D_2O$) δ 172.9, 52.7, 36.1, 35.5, 32.0, 31.7, 24.6, 24.4.

MS (ESI): calc. for $[M+H]^+$: 158.12, obs.: 158.2.

#### Cyclohepthylmethanol

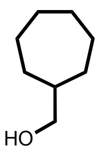

Cycloheptanecarboxylic acid (1.01 g, 7.03 mmol, 1.0 eq.) was dissolved in dry THF (20 mL) under a nitrogen atmosphere, and the solution was cooled to 0 °C. $LiAlH_4$ (2.4 M in THF, 8.8 mL, 21 mmol, 3.0 eq.) was added dropwise, and the reaction was stirred at 0 °C for 1 h, followed by 1.5 h at room temperature. The reaction mixture was cooled to 0 °C, and water (2 mL), aq. NaOH (15%, 2 mL), and water (4 mL) were added. The slurry was stirred for 10 min and filtered through Celite. The filter cake was washed with EtOAc (3x), and the combined organic phases were washed with water and brine. The organic phase was dried over $Na_2SO_4$, and the supernatant was collected by filtration. The solvent was removed under reduced pressure to yield cyclohepthylmethanol (900 mg, 7.03 mmol, quant.) as a colorless oil.

$^1$H NMR (400 MHz, $CDCl_3$) δ 3.42 (d, $^3J_{HH}$ = 6.6 Hz, 2H), 1.80 – 1.38 (m, 11H), 1.25 – 1.13 (m, 2H).

**Cyclohepthylmethylbromide**

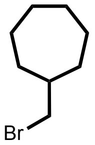

Cyclohepthylmethanol (900 mg, 7.03 mmol, 1.0 eq.) and $CBr_4$ (3.04 g, 9.17 mmol, 1.3 eq.) were dissolved in dry DCM (13 mL) under a nitrogen atmosphere, and the solution was cooled to 0 °C. A solution of $PPh_3$ (2.41 g, 9.17 mmol, 1.3 eq.) in dry DCM (13 mL) was added dropwise, and the mixture was stirred at 4 °C for 3 h. The solvent was evaporated under reduced pressure, n-pentane (30 mL) was added, and the suspension was sonicated for 5 min. The flask was chilled in a freezer for 30 min, and the suspension was filtered. The precipitate was washed with chilled n-pentane, and the precipitation was once more worked up by a cycle of suspension, sonication, chilling, and filtering. Both filtrates were combined, the solvent was removed under reduced pressure to yield cycloheptylmethylbromide as a yellowish oil (1.08 g, 5.68 mmol, 74%). There was still triphenylphosphine oxide present, but it was used without further purification for the next step.

$^1$H NMR (400 MHz, $CDCl_3$) δ 3.32 (d, $^3J_{HH}$ = 5.9 Hz, 2H), 1.92 – 1.22 (m, 13H). With triphenylphosphine contamination in the aromatic region.

**Diethyl 2-acetamido-2-(cycloheptylmethyl)malonate**

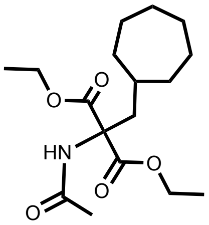

Diethyl acetamidomalonate (1.03 g, 4.76 mmol, 1.0 eq.) was dissolved in DMSO (2 mL) under a nitrogen atmosphere. $Cs_2CO_3$ (2.33 g, 7.14 g, 1.5 eq.) and cycloheptylmethylbromide (1.00 g, 5.23 mmol, 1.1 eq.) in DMSO (3 mL) were added and the suspension was stirred at 60 °C for 17 h. The reaction was allowed to reach room temperature, and ice-cold water (60 mL) was added. The mixture was extracted with EtOAc (2 x 60 mL), and the combined organic phases were washed with brine (100 mL). The organic phase was dried over $Na_2SO_4$, and the supernatant was collected by filtration. The solvent was removed under reduced pressure to give an orange oil. The crude product was purified by automated flash column chromatography (25 g $Si_2O$, 15% EtOAc in cyclohexane to 100% EtOAc) to yield an orange solid. Although the product still contained minor impurities, the desired compound peaks could be identified, and the crude material was taken for the next step.

$^1$H NMR (400 MHz, $CDCl_3$) δ 4.23 (q, $^3J_{HH}$ = 7.1 Hz, 4H), 2.33 (d, $^3J_{HH}$ = 6.0 Hz, 2H), 2.03 (s, 3H), 1.65 – 1.16 (m, 13H), 1.25 (t, $^3J_{HH}$ = 7.1 Hz, 6H).

**Cycloheptylalanine hydrochloride (C7a)**

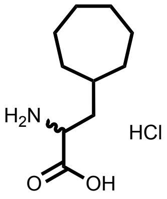

Crude diethyl 2-acetamido-2-(cycloheptylmethyl)malonate (approx. 570 mg, 1.74 mmol, 1.0 eq.) was dissolved in HCl (4 M in dioxane, 10 mL) and stirred at 100 °C for 16 h. Starting material was still observed by LC-MS. Thus, additional aq. HCl (4 M, 10 mL) was added. The reaction mixture was stirred at 100 °C for another 24 h, and the solvents were evaporated under reduced pressure to yield cycloheptylalanine hydrochloride (145 mg, 654 µmol, 13% over 2 steps) as a brownish solid.

$^{1}H$ NMR (400 MHz, $D_2O$) δ 4.01 (dd, $^{3}J_{HH}$ = 8.4, 5.8 Hz, 1H), 1.96 – 1.86 (m, 1H), 1.80 – 1.38 (m, 12H), 1.33 – 1.20 (m, 2H).

$^{13}C$ NMR (101 MHz, $D_2O$) δ 173.3, 51.7, 38.2, 34.6, 34.0, 32.9, 27.9, 27.9, 25.5, 25.3.

MS (ESI): calc. for $[M+H]^{+}$: 186.15, obs.: 186.2.

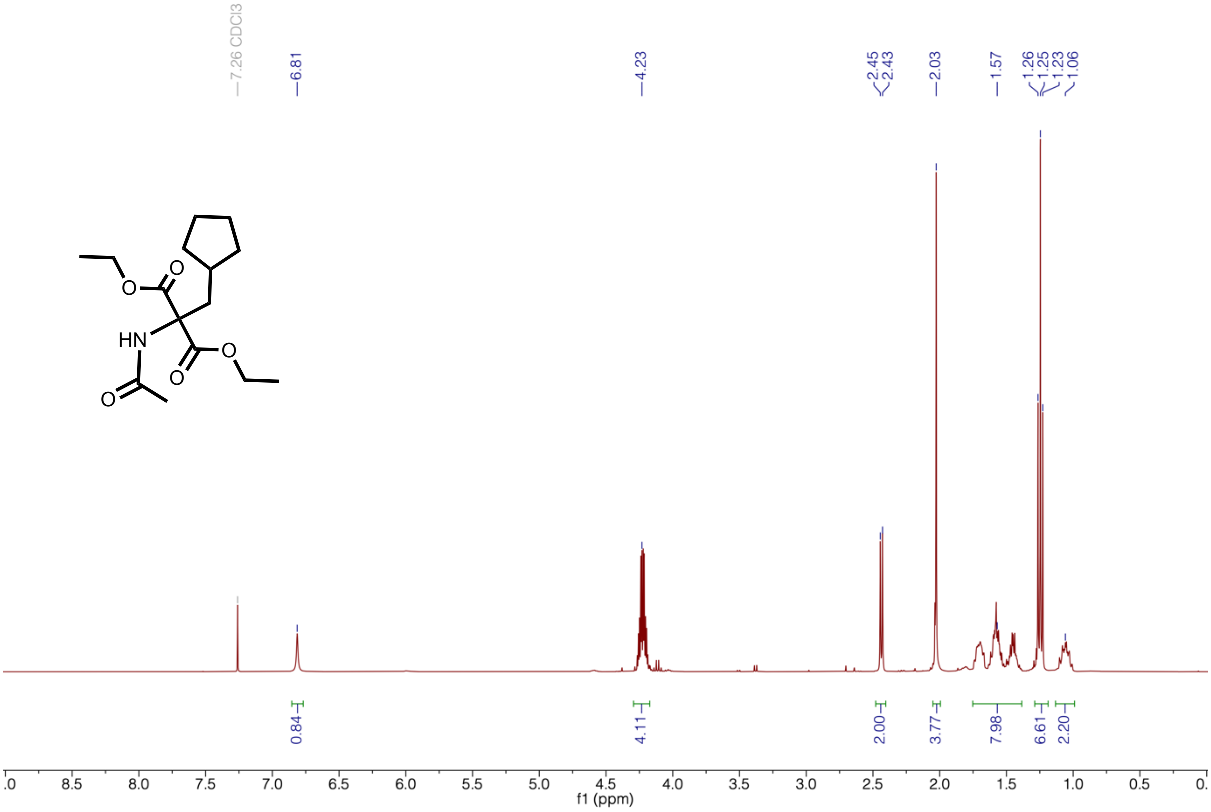

**Supplementary Figure 9 | $^{1}$H-NMR of Diethyl 2-acetamido-2-(cyclopentylmethyl) malonate.**

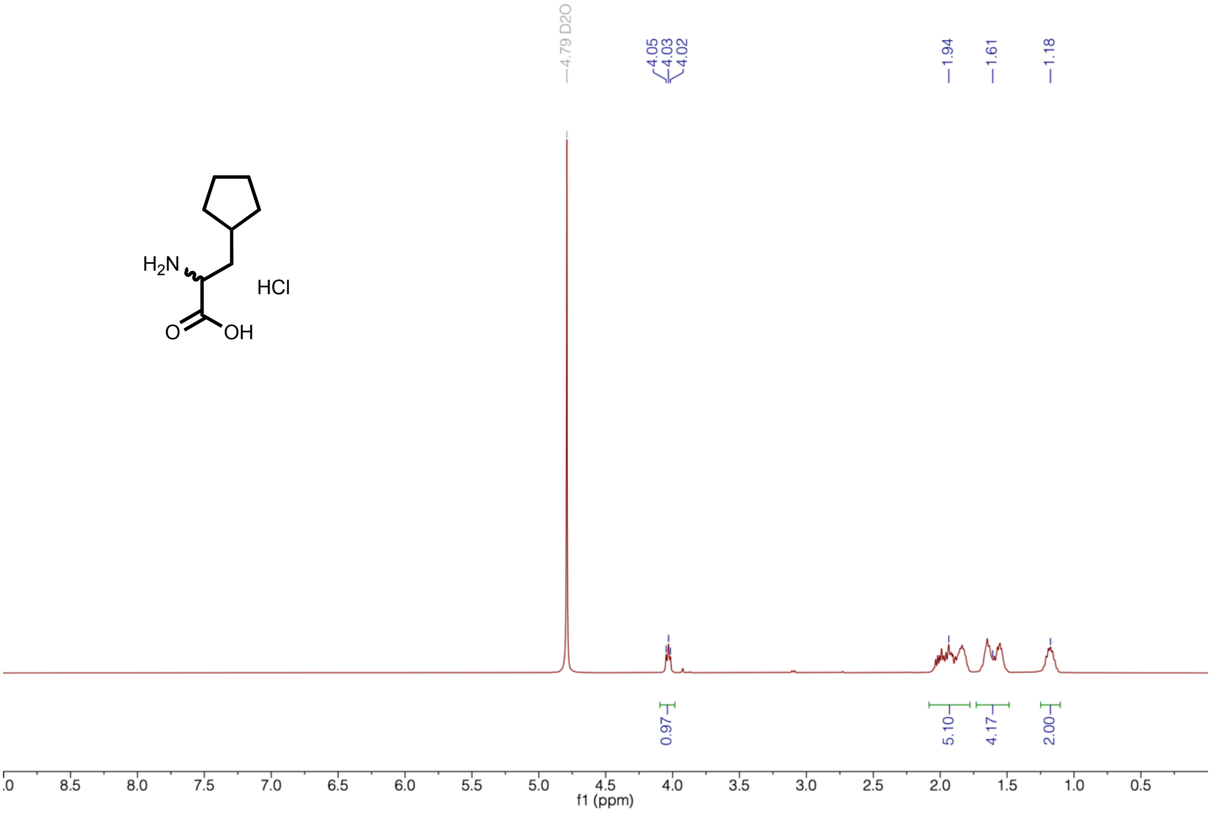

**Supplementary Figure 10 | $^{1}$H-NMR of C5a**

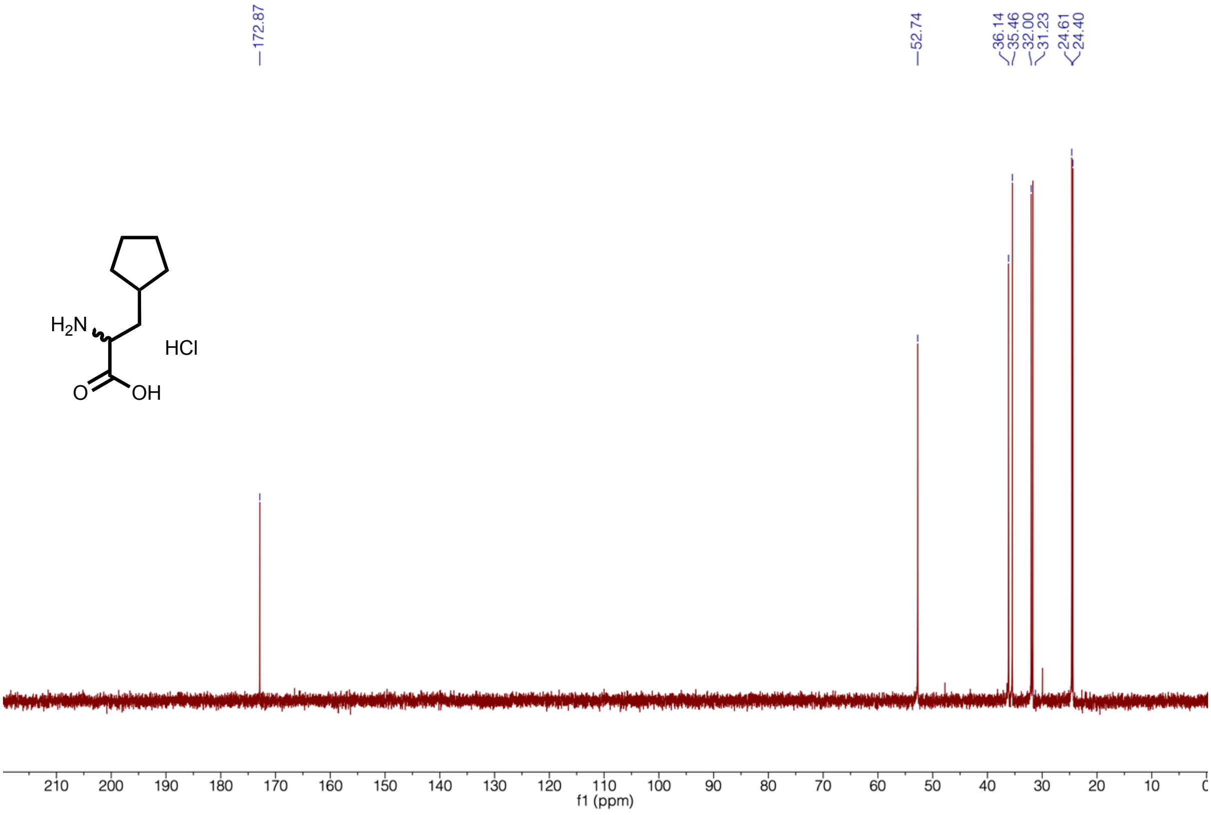

**Supplementary Figure 11 | $^{13}$C-NMR of C5a**

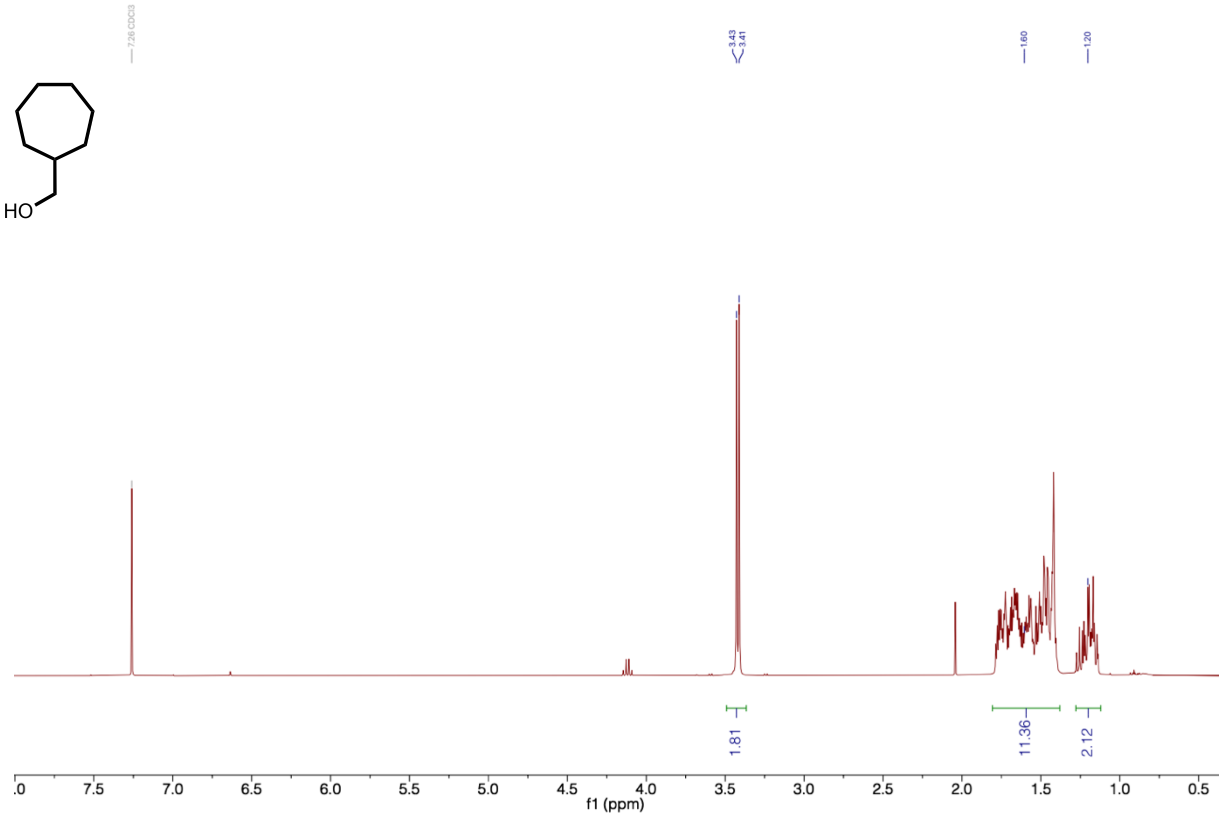

**Supplementary Figure 12 | [1]H-NMR of cyclohepthylmethanol**

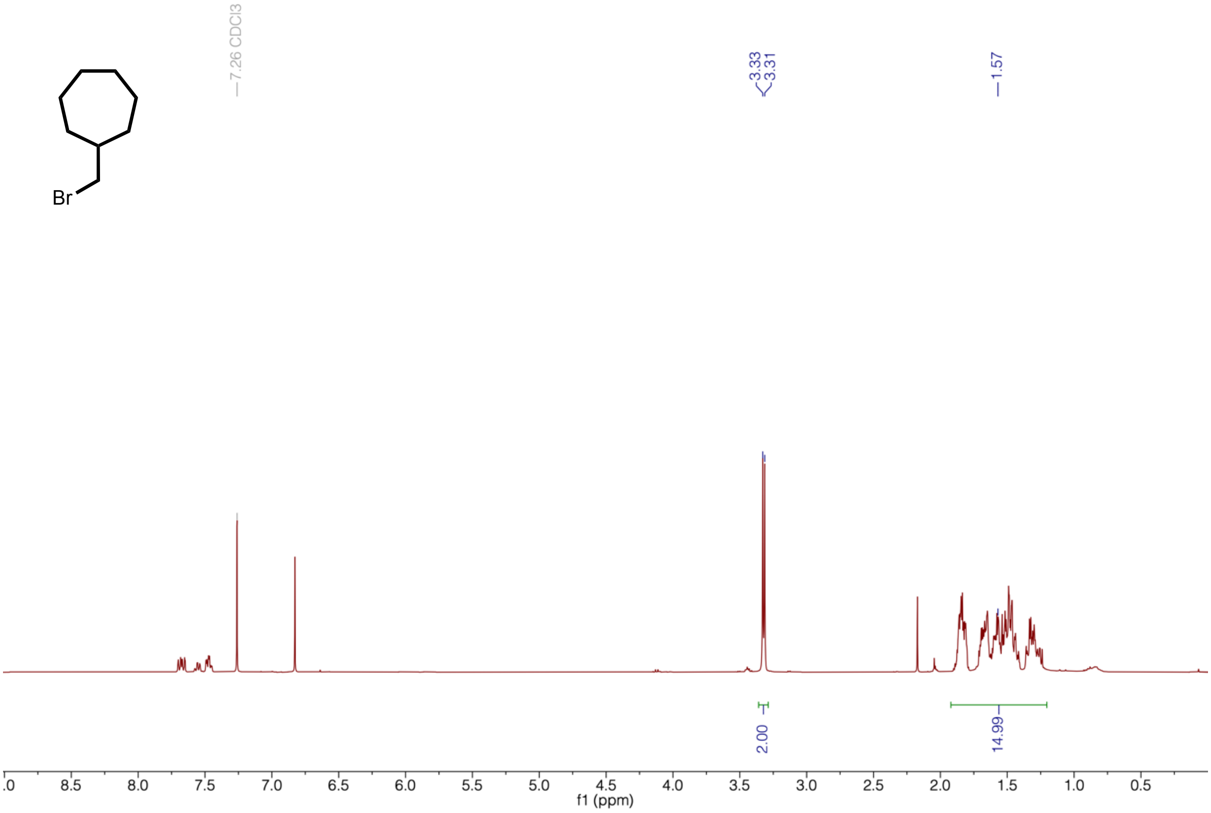

**Supplementary Figure 13 | [1]H-NMR of crude cyclohepthylmethylbromide**

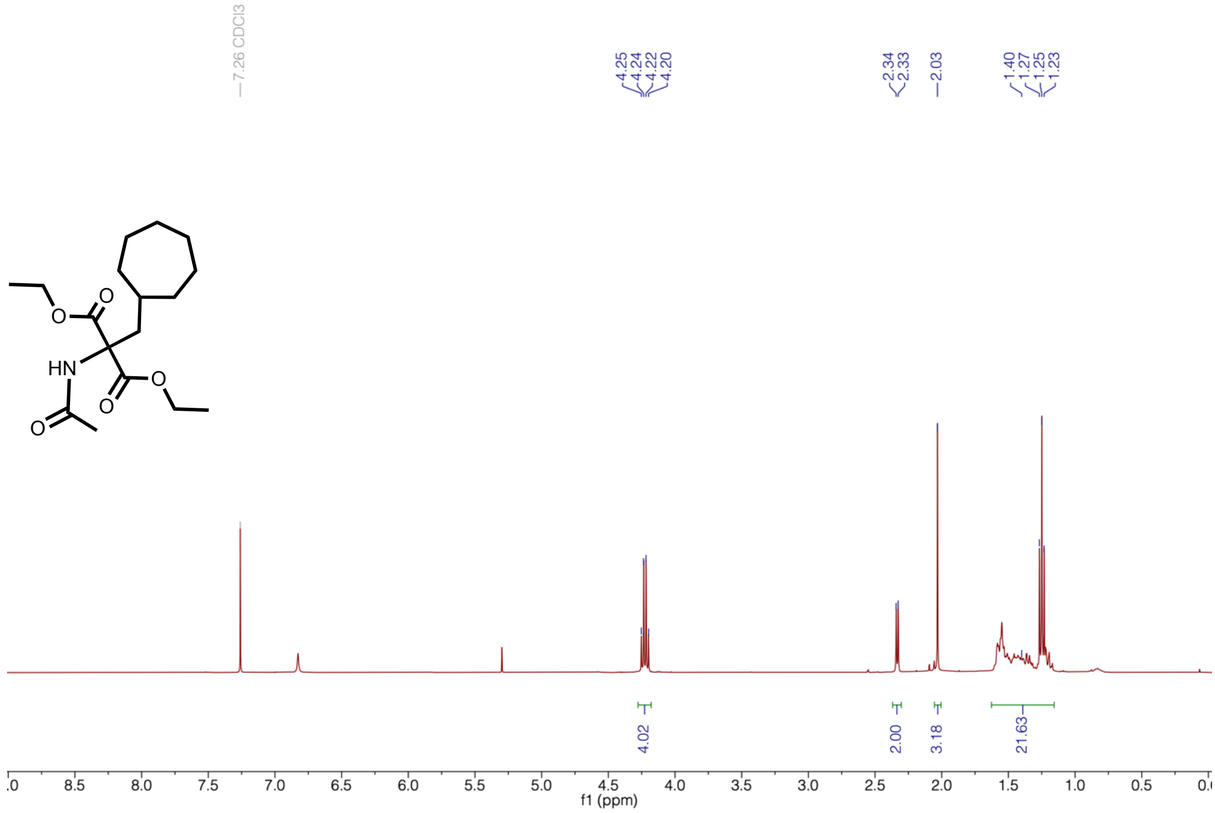

**Supplementary Figure 14 | $^1$H-NMR of diethyl 2-acetamido-2-(cycloheptylmethyl) malonate**

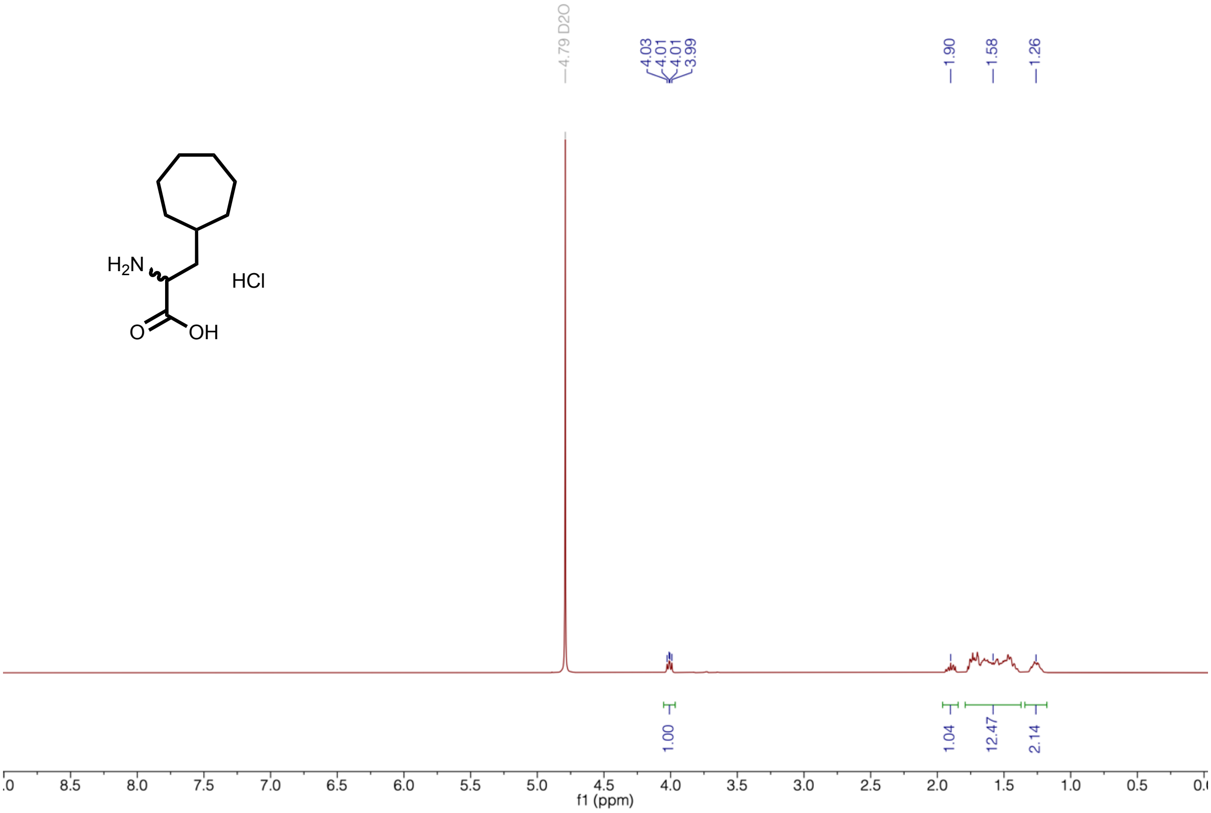

**Supplementary Figure 15 | ¹H-NMR of C7a**

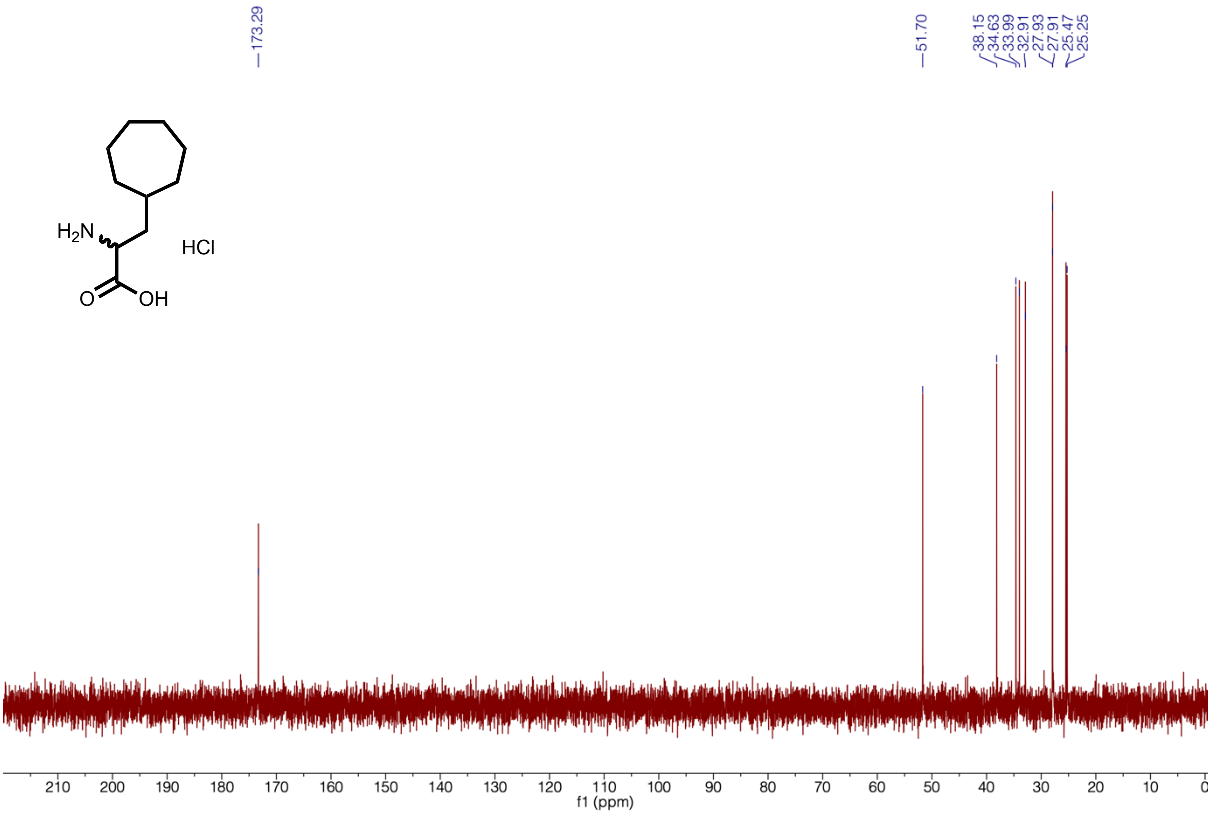

**Supplementary Figure 16 | ¹³C-NMR of C7a**

### III. Kinetic parameters

**Supplementary Table 1 | Summary parameters for *Sc*SLAC variants.**

| **Variant** | **$E°'_{T1Cu}$ (mV)** | **$\Delta E°'_{T1Cu}$ (mV)** | **$k_{cat}$ ($s^{-1}$)** | **$K_m$ (mM)** | **$k_{cat}/K_m$ ($s^{-1}$ $mM^{-1}$)** |
|---|---|---|---|---|---|
| M298 | 359 ± 8 | -- | 1.6(8) | 0.96(5) | 1.6(2) |
| M298**C5a** | 452 ± 12 | +93 | 3.1(1) | 1.27(2) | 2.5(1) |
| M298**C6a** | 475 ± 10 | +116 | 3.0(2) | 1.05(5) | 2.8(1) |
| M298F | 353 ± 5 | -6 | 0.19(3) | 0.4(1) | 0.46(4) |
| M298L | <350 | -- | 0.32(2) | 1.19(7) | 0.27(1) |

Entries represent mean and standard deviation.

### IV. Crystallography

**Supplementary Table 2 | Crystallography: data collection and refinement statistics.**

| | **ScSLAC M298C6a** |
|---|---|
| | *native* |
| **Data collection** | |
| Wavelength (Å) | 0.9677 |
| Space group | P 43 3 2 |
| *a, b, c* (Å) | 178.76, 178.76, 178.76 |
| α, β, γ (°) | 90, 90, 90 |
| Resolution (Å) * | 63.20 – 3.49 (3.59 – 3.49) |
| $R_{meas}$* | 0.424 (1.467) |
| $R_{merge}$* | 0.391 (1.370) |
| Mean I/σI* | 5.4 (1.7) |
| Completeness ellipsoidal (%)* | 96.5 (62.7) |
| Multiplicity* | 6.6 (7.8) |
| CC1/2* | 0.961 (0.511) |
| **Refinement** | |
| Resolution (Å) | 63.20 – 3.49 |
| Total reflections* | 82690 (4830) |
| Total unique* | 12461 (623) |
| $R_{work}$# | 0.1874 (0.2527) |
| $R_{free}$# | 0.2305 (0.2803) |
| Number of non-hydrogen atoms | 2149 |
| macromolecules | 2145 |
| ligands | 4 |
| solvent | 0 |
| Protein residues | 277 |
| RMS deviations (bonds)# | 0.011 |
| RMS deviations (angles)# | 1.47 |
| Ramachandran favored (%)# | 97.79 |
| Ramachandran allowed (%)# | 2.21 |
| Ramachandran outliers (%)# | 0.00 |
| Rotatmer outliers (%)# | 0.00 |
| Clashscore# | 3.83 |
| Average B-factor# | 66.54 |
| macromolecules | 66.49 |
| ligands | 93.71 |
| Molprobity Score | **1.22** |
| PDB | **9HU7** |

#as reported by phenix.table_one and phenix.model_vs_data
*as reported by autoPROC and phenix.table_one
